# A comprehensive conceptual and computational dynamics framework for Autonomous Regeneration Systems

**DOI:** 10.1101/820613

**Authors:** Tran Nguyen Minh-Thai, Sandhya Samarasinghe, Michael Levin

**Affiliations:** Complex Systems, Big Data and Informatics Initiative (CSBII), Lincoln University, New Zealand; Tufts Center for Regenerative and Developmental Biology, Tufts University, Boston, USA

## Abstract

This paper presents a new conceptual and computational dynamics framework for damage detection and regeneration in multicellular structures similar to living animals. The model uniquely achieves complete and accurate regeneration from any damage anywhere in the system. We demonstrated the efficacy of the proposed framework on an artificial organism consisting of three tissue structures corresponding to the head, body and tail of a worm. Each structure consists of a stem cell surrounded by a tissue of differentiated cells. We represent a tissue as an Auto-Associative Neural Network (AANN) with local interactions and stem cells as a self-repair network with long-range interactions. We also propose another new concept, Information Field which is a mathematical abstraction over traditional components of tissues, to keep minimum pattern information of the tissue structures to be accessed by stem cells in extreme cases of damage. Through entropy, a measure of communication between a stem cell and differentiated cells, stem cells monitor the tissue pattern integrity, violation of which triggers damage detection and tissue repair. Stem cell network monitors its state and invokes stem cell repair in the case of stem cell damage. The model accomplishes regeneration at two levels: In the first level, damaged tissues with intact stem cells regenerate themselves. Here, stem cell identifies entropy change and finds the damage and regenerates the tissue in collaboration with the AANN. In the second level, involving missing whole tissues and stem cells, the remaining stem cell(s) access the information field to restore the stem cell network and regenerate missing tissues. In the case of partial tissue damage with missing stem cells, the two levels collaborate to accurately restore the stem cell network and tissues. This comprehensive hypothetical framework offers a new way to conceptualise regeneration for better understanding the regeneration processes in living systems. It could also be useful in biology for regenerative medicine and in engineering for building self-repairing biobots.

## 1. Regeneration in Biology – inspiration for self-repair biological and artificial machines

Regeneration of complex structures is a wonderful phenomenon in nature and it typically involves the action of recovery of body parts lost by amputation as well as dying cells. It is an ancient process, but it is not completely understood (Maden, 1992; Needham, 1961). The capacity for regeneration varies greatly among animals; for example, snails can regrow their heads, axolotls can regenerate their limbs, zebrafish can reproduce new hearts (Rabinowitz et al., 2017) and Hydra and planarian can recover the whole animal from a tiny piece of their bodies (Frei et al., 2013; Major and Poss, 2007). Likewise, plants also have a high capacity to regenerate which has been used for producing new plants by cutting and grafting. Regeneration in mammals is almost completely limited except for deer antlers and liver. Humans, for example, can recover from wounds and surgeries and make new skin and blood cells (Wilgus, 2007).

Most natural structures can regenerate to a small or large extent, which accordingly contributes to their robustness and resilience. However, little is known about the mechanisms of regeneration of biological organisms. Regeneration is achieved through a process where specific cells, such as stem cells, divide to produce new (differentiated) cells to regenerate the original pattern and stop when it is complete. In the process, a large number of tissue cells interact with each other to accomplish regeneration, but it is still unclear how these interactions lead to the exact recovery of the lost body form. Although the molecular mechanisms required for regeneration are being discovered, the algorithms sufficient for the regeneration of complex anatomical structures (not merely cell differentiation) represent a significant knowledge gap that holds back evolutionary developmental biology and regenerative medicine. The focus of this study is to contribute towards the development of a *conceptual framework* for a complete regeneration system for a simple organism like planarian. It is hoped that this could also be a platform for implementing self-repairing bio-robots and artificial robots and contribute to bio-inspired computing.

Planaria are well known for the ability to completely regenerate the original form. It is a flatworm living in both freshwater and saltwater. A tiny piece as small as 1/279th of a planarian can regenerate into a new animal within approximately two weeks (Morgan, 1898). Planaria produce new tissues from the unique stem cells known as neoblasts to restore damage regions (Wagner, Wang, & Reddien, 2011). These stem cells can migrate through the whole organism into the blastema to regenerate missing parts. An interesting aspect of this regeneration is that a tiny piece develops into a small worm that grows into the full form by concurrent regeneration of body parts. Another exciting fact is that a small trunk fragment of a planarian after cutting the head and the tail always regenerates the head and tail in the correct orientation as in the original planarian. It indicates that the remaining tissue identifies the anterior-posterior (abbreviated as A/P) polarity that helps new parts grow in the correct positions. Planaria also have another polarity dorsal-ventral (abbreviated as D/V) or top-bottom polarity. A/P polarity in planarian is similar to a bar magnet in that a new bar magnet after being cut transversely also restores the original North-South polarity (Lobo, Beane, & Levin, 2012). However, regeneration in planaria is a mysterious mechanism that involves complex interactions and communications between cells at the organismal level to maintain complex shapes after amputation. How polarity is established in pre-existing tissue during regeneration is also still unclear. Therefore, as a paradigmatic example of regeneration, the discovery of the mechanism (molecular and algorithmic) of planarian regeneration has always been fascinating to researchers, particularly those in Developmental and Regenerative Biology, as many basic issues in pattern formation and regeneration are still open questions. This paper contributes to the algorithmic mechanisms of regeneration.

Some researchers have introduced algorithmic models of regeneration that have focused on signal communication between stem cells and differentiated cells in a tissue (Bessonov et al., 2015; De, Chakravarthy, & Levin, 2017; Ferreira, Smiley, Scheutz, & Levin, 2016; Tosenberger et al., 2015). These models have proposed rules for regenerating a given morphology and the existence of a coordinate system that guides the patterning of the organism. However, none of the previous models can explain how a stem cell can regenerate a whole body after amputation and they require stem cells and other cells to remember too much information. Further, they have difficulty stopping the regeneration process at the right time leading to excessive growth and pruning. These models also carry a large computational burden due to excessive communications. In our previous study (Minh-Thai, Aryal, Samarasinghe, & Levin, 2018), we proposed an autonomous self-repair system for regenerating a simple tissue after damage based on effective collaboration between a stem cell and differentiated cells in the tissue using a neural network framework. The model achieves complete and accurate regeneration with minimum computation. Therefore, as a high-level conceptual system, this model offers promising extensions to complex pattern regeneration that we aim to explore in this study.

In recent years, there has also been an increasing interest in self-repair systems in robotics (Arbuckle & Requicha, 2010; Rubenstein, Sai, Chuong, & Shen, 2009). Some of these systems in a simple way use the concept of stem cells (Rubenstein et al., 2009). This field is also in its infancy and much can be done to develop interesting robotic self-repair systems that mimic biological regeneration for various purposes including synthetic biology and regenerative medicine.

Thus, although some progress has been made in understanding and modelling regeneration, much needs to be done to address the fundamental questions of regeneration and develop effective algorithms to mimic the regeneration process. These fundamental questions in general characterise the philosophical, biological, and algorithmic nature of regeneration of living tissues. The most fundamental question, in the case of planarian regeneration as a paradigmatic example, is – how does a small portion of the body know the whole pattern? This gives rise to a whole set of other questions, such as, is pattern information stored somewhere?; If so, where is it stored, how is it retrieved and what mechanisms are used to implement it during regeneration? What oversees or drives regeneration?; How does the biological system respond to such a pattern? What is the minimum size of the body fragment sufficient for complete regeneration? Alternatively, if it is a case of regeneration without a stored pattern in existence, then, where does patterning information come from, what drives regeneration, and how does it know when to stop regeneration to avoid excessive growth? At a more working level – what senses and communicates the damage and what receives this information and in what form? And what computational paradigms are involved in this communication and regeneration? These and many are as yet unanswered questions in biological regeneration.

Advances in evolutionary developmental biology and applied regenerative medicine require a quantitative understanding of the control systems of regeneration, which will enable design of interventions for rational control of system-level anatomical properties. Here, we propose a study of computational dynamics that could guide regenerative mechanisms in evolved or artificial (synthetic) systems. The research focuses on some of the above questions through the extension of our previous autonomous tissue self-repair system in (Minh-Thai et al., 2018) to a regeneration system for a complete organism. Our goal is to develop a computational framework that conceptually mimics planarian regeneration in some important aspects: regeneration of the whole pattern even from a tiny fragment, concurrent regeneration of all affected tissues, complete and accurate regeneration without overgrowth or pruning and minimum computational burden. As a generic framework, it could also benefit synthetic biology, and promote advanced self-regenerating bio-robots and artificial robots.

## 2. Regeneration as a biological computing process

(Bessonov et al., 2015) presented a conceptual framework for regeneration based on signal distribution and cellular memory. They assumed that cells communicate with each other using signals that are produced by individual cells and released into the surrounding space. These signals decay with distance as a function f(d)=1/d^n^. Each cell keeps the value of the total signal (old signal) that it received from all the other cells. After amputation, each cell computes a new value of the total signal (current signal) based on communication received from the remaining cells. The difference between the old signal and the current signal stimulates the regeneration of a correct cell structure after damage using some rules of regeneration. Regeneration of this model depends on the value of a parameter *n* in the decay function (f(d)=1/d^n^) and on the size of the structure. The model allows correct regeneration of a small and convex structure when the value of *n* is two. This model works incorrectly for a large organism as it incorrectly identifies missing cells at a large distance because the signal decays quickly. The model also requires too much computation for calculating the total signal frequently and compare the current total value with the old value to determine the damage. There is also too much communication between cells as a cell can communicate with all other cells. Each cell also needs to keep its total value of signals for regeneration. In this model, every cell can divide into new cells even if it is not a stem cell which may not be very realistic in biological regeneration.

Another model of regeneration was proposed by (Tosenberger et al., 2015). It described the development and regeneration of a structure where a global submodel presents the restoration of tissue centres (e.g., stem cells), while the local model produces tissues around the stem cell. In the global model, they alter the position of stem cells and make them return to the initial positions. In the first case of this model, they assume that all stem cells produce the same type of signal. However, in this mode, there is not enough information to restore correct stem cell positions. In the second case, each stem cell produces its own signal which is different from signals from the other stem cells. In this case, the model can restore the exact initial configuration of the stem cell structure when the positions of stem cells are altered. The local model describes how a tissue grows around stem cells. The tissue consists of a stem cell at the centre surrounded by differentiated cells. The model assumes that the stem cell produces a decaying signal which spreads in the tissue space. Every differentiated cell receives this signal and then compares with a predefined threshold to decide whether to kill itself or stay. This model allows correct regeneration of a circular tissue with minimum assumptions that can apply to bibots. It is more bio-realistic than the previous model in that only stem cells can divide. However, in the global model, the assumption is that stem cells do not get damaged; therefore, it cannot replace them if they are damaged. Also the model restores unsuccessfully in the case where a stem cell moves in a diametrically opposite direction. In the local model, regeneration works correctly only for small tissues due to signal decay. Also, stem cells divide continuously to keep the correct shape but cells have to be killed unnecessarily to attain the correct shape.

An agent-based model is proposed in (Ferreira et al., 2016) to discover a 3D structure of stem cells and self-repair when a part is cut. The mechanism discovers the whole structure and repairs it from damage or cell death. Each cell creates, sends and receives messages (or packets) to other cells. The information in the packets is used to form a structure as well as detect and repair any dead/missing cells. They improved the performance of the model by applying noise to the direction and distance of packets (Ferreira, Scheutz, & Levin, 2017b). Another improvement is presented in (Ferreira, Scheutz, & Levin, 2017a) where they reduced the number of packets by using two types of cells: neoblasts (stem cells) and differentiated cells. Packets are only produced by the neoblasts and the differentiated cells only relay such packets. However, too many packets need to be kept in the stem cells as well as too much communication between cells takes place when they maintain and regenerate the system.

Another computational approach using interactions between a neural network (nervous system) and non-neural (body) cells to help an organism grow and recover its initial form from damage is presented in (De et al., 2017). Non-neural cells communicate with the neural cells as well as each other in the neighbourhood to modulate four biological variables. Neural cells provide patterning cues to body cells in terms of A/P polarity based on bioelectric signals. This is by far the most biologically realistic model with a reasonable regeneration success (55-80%). Limitations of the model are that excessive pruning is required during regeneration and the difficulty in regenerating the exact final form. A genetic algorithm model proposed by (Gerlee, Basanta, & Anderson, 2011) evolves and maintains a 3D cellular tissue structure. The algorithm shows that one cell evolves into a network that is similar to cells organized in some of the tissues of the body. However, a predefined area needs to be defined for the network to grow; otherwise, the network grows indefinitely. Thus, some interesting algorithmic ideas of regeneration have been tested with varying degrees of success; however, most of the past regeneration models are either computationally expensive due to extensive communications and/or the type of algorithms or not biologically realistic.

One of the issues in planarian regeneration is how the damaged tissue re-establishes the polarity during regeneration (Morgan, 1901). Previous studies have reported that Wnt and BMP signalling pathways are necessary for polarity patterning (Little & Mullins, 2006; Niehrs, 2004). These signals affect polarity while planarian regenerates lost body parts. However, recently, Pietak et al. (2019) have shown that neural directionality can set the biochemical gradients that determines polarity.

Incidentally, regeneration has been popular in robotics as well. A swarm of robots has been programmed to construct and self-repair two-dimensional structures (Arbuckle & Requicha, 2010; Rubenstein et al., 2009). Inspired by biological stem cells, robotic stem cells are robots that have the capacity to self-organise and self-heal a damaged structure. Robotic stem cells are each programmed to remember a copy of the whole structure. They reconstruct the same pattern when the original structure is damaged or there is a difference between the desired and the current shape. When robots suffer damage, the remaining robots may reconfigure and reorganize the same pattern but on a smaller scale (smaller size) and continue to function. This process to some extent is analogous to regeneration in biology. From a robotic perspective, apart from not achieving the scale of the original pattern, how they can learn to form new patterns could be worth further research such as reorganising into new structural patterns that in a way mimics biological evolution of forms that are resilient.

In our work (Minh-Thai et al., 2018) mentioned previously, we presented an autonomous tissue self-repair system for damage detection and regeneration inspired by biological regeneration. The model considers a circular tissue consisting of a stem cell surrounded by thousands of differentiated cells. The tissue is considered as a system that is aware of its own pattern integrity (self-aware) and reacts to damages to efficiently restore it. To achieve this, the tissue is represented as a self-aware neural network system. Specifically, it is a bi-directional graph in the form of an Auto-Associative Neural Network (AANN) consisting of neurons (differentiated cells) that communicate with their immediate neighbours. Further, the stem cell communicates with the neurons to maintain tissue integrity and direct the repair process after damage and the AANN assists the stem cell restore the pattern.

This self-repair system consists of two sub-models– Global sensing (GS) and Local sensing (LS). Global sensing uses entropy as a measure of tissue integrity maintained by the stem cell through global communication with the differentiated cells in a way that the change in the system state due to damage is recognized by the stem cell by a change in entropy that enables it to detect the overall damage region in the tissue. Then, local sensing is triggered in the corresponding region of the AANN to identify the exact damage location using local communications among differentiated cells, following which the stem cell migrates to the damage location and regenerates the damaged tissue. This two level communication with focused activation of only the required region of the AANN greatly *minimises the amount of computation* needed for regeneration; also, the model *achieves complete regeneration* from damage of any size almost anywhere in the tissue – thus it addresses the two main limitations of the previous models. The system fully recovers from a wide variety of damages as long as at least one string of cells from the stem cell to the border remains to provide pattern information during regeneration. Although this covers a large spectrum of damages that the system can handle, having to retain pattern information in the tissue, that is not possible for extreme damages where the whole outer region of the tissue goes missing, is still a limitation of this model. Also, it has only been applied to a simple tissue of circular shape.

Some of the above-mentioned important advantages of our system over the existing methods provide the scope for improving and extending it to other tissue patterns and multiple tissue systems to represent regeneration in simple organisms. This is the aim of the current study. This brings out some additional fundamental questions about the mechanism of planarian regeneration and provides the opportunity to offer hypothetical solutions. For example, there are two basic mechanisms of regeneration in planaria: when a local tissue is damaged, stem cells sense and migrate to the damaged area to correctly repair the damage (our previous model has covered this). When any part with stem cells is cut off, the remaining stem cells sense the damage, move to the blastema, produce new stem cells and regenerate the missing tissues concurrently; we aim to incorporate this into our model. The additional questions related to these two mechanisms are: how does a system know the pattern of its various tissues (head, body, tail etc.)?; how are various tissue systems in an organism maintained and coordinated concurrently in regeneration? how do stem cells communicate with each other? how do stem cells and tissue cells in a multiple tissue system collectively implement regeneration?. These questions allow great scope for models that are not only biologically realistic but also computationally efficient. This research attempts to offer hypotheses to answer these questions in order to make improvements towards efficient regeneration systems that completely and correctly regenerate their form that in a broad sense resembles biological systems.

In this paper, we extend our previous tissue self-repair model to other tissue shapes and organise them into a conceptual regeneration system for a whole organism where stem cells detect tissue as well as stem cell damage and regenerate new stem cells and tissues to recover the whole organism after damage. When a damage happens anywhere in the organism, it efficiently senses the nature of the damage, whether a tissue or stem cell damage, detects the location and regenerates the missing stem cells, whole tissues, or tissue parts to bring the system to its exact original state.

In the next section, we introduce the overall concept of the new system and demonstrate its efficacy in recovering from all possible types of damage.

## 3. A new conceptual and computational dynamics framework for whole system regeneration

This section develops the proposed whole system regeneration model. Here we introduce some new concepts to mimic planarian regeneration. We hypothesise that a system that is able to regenerate its original pattern from any tiny fragment anywhere in the body after a damage must be extremely versatile and robust, and therefore, it possibly uses a high-level and flexible algorithm, as opposed to extensive micromanagement, to regenerate itself. Further, since a very tiny fragment can regenerate the whole pattern, we hypothesise that the pattern information must either be stored everywhere in the organism or be accessed by any part of the organism from an as yet unknown space which we call a virtual information field surrounding the organism. To represent these ideas, we introduce two new concepts: smart stem cells with virtual information fields that carry minimum pattern information relevant to the corresponding tissues (head, tail etc.) (tissue memory) so that any damaged tissue can be repaired at the tissue level; and a stem cell network, corresponding to the whole organism, that creates a shared information field (collective memory) by combining individual stem cell information fields so that any missing stem cells can be identified and regenerated for recovering the whole pattern from any small or large scale damage.

Specifically, we first extend our original circular tissue model to introduce a smart stem cell with a virtual information field that holds the minimum pattern information to enhance the capability of the original stem cell. This completely releases the system from the constraint of having to retain minimum pattern information in the damaged tissue. We then extend the model to more complex (triangular and rectangular) tissue shapes each with a smart stem cell and a virtual information field that holds the minimum pattern information for the corresponding tissues. These tissue shapes are then combined to form a simple virtual organism consisting of three parts – head, body and tail-each with a smart stem cell surrounded by a large number of differentiated cells. The organism thus contains smart stem cells, differentiated cells and a combined virtual field that holds the minimum pattern for the whole organism (collective memory). Stem cells form a network that monitors the state of the whole organism and recovers itself from any damage. For this, we develop a self-repair stem cell network model and explore three methods for its implementation: Automata, Neural Networks and Decision Trees. Differentiated cells form an AANN with local neighbourhood communication.

In this new system, the smart *stem cell network repair model* and individual *tissue repair models* collaborate seamlessly to maintain the integrity of the system under any damage condition by autonomously regenerating missing stem cells and damaged tissues. In most cases of tissue damage, the system would regenerate tissues without tapping into their individual information fields. However, in extreme cases of tissue damage that create open-ended boundaries, a stem cell accesses minimum pattern information from the field, thereby making it possible to recover from any tissue damage. In the case of stem cell damage, the stem cell network taps into the collective field to regenerate new stem cells to continue regeneration. The combined system mimics some of the main features of planarian regeneration, including regeneration of the whole pattern from any tiny fragment as long as at least one stem cell (regardless of which tissue) remains after damage; complete and accurate regeneration from minor damages to individual and/or multiple tissues or severe damage incurring loss of whole body parts (head, body or tail). Thus, the proposed system is robust, versatile and flexible with complete regeneration ability achieved with minimum computation. Below we describe the details of the new system starting with an overview of our original tissue repair model followed by the proposed extensions.

### 3.1. Base model: Autonomous self-repair system – circular tissue

Our original tissue repair model in (Minh-Thai et al., 2018) describes regeneration of a virtual tissue which consists of a stem cell surrounded by a thousand differentiated cells in a 2-D circular tissue. Each cell has a pair of polar coordinates (*radius, θ*) (Fig. 1) with a small noise added from Brownian motion to represent uncertainty in its position or any slight movement thereof. The stem cell can divide itself into two new cells (a stem cell and a differentiated cell) to replace dying or missing cells in the tissue. The tissue is represented by an Auto-Associative Neural Network (AANN) where each differentiated cell is a perceptron (threshold neuron) with connections to the immediate neighbours (i.e., locally recurrent network). This represents a self-aware system that autonomously preserves itself, that is achieved through two types of communication: Global communication that represents the communication between the stem cell and all differentiated cells for maintaining the overall tissue integrity; and local communication that describes the form of communication between immediate neighbour cells in the AANN for local management of the pattern and assisting in repair.

**Fig. 1.**
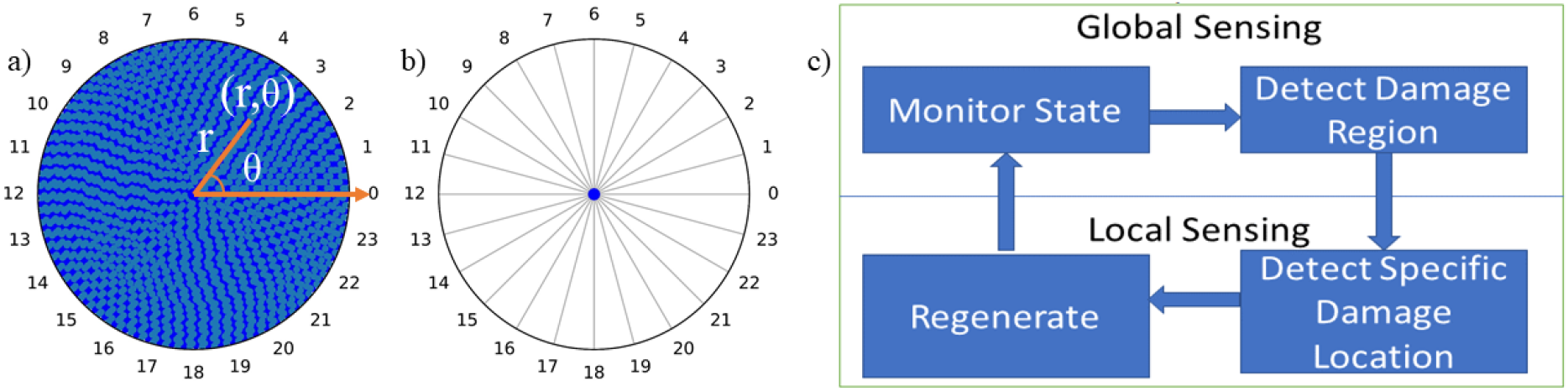
a) A stem cell at the center surrounded by differentiated cells, b) Tissue pattern divided into 24 regions to help specify global damage regions by Global Sensing, and c) Functional aspects of the tissue self-repair framework

#### 3.1.1 Operation of the base model

Fig.1 shows the system with its two global sensing (GS) and local sensing (LS) sub-models. Global sensing uses entropy and change in entropy for sensing changing states. In a real biological system, damage can spill electrolytes and signalling molecules into the extra cellular matrix (ECM), that can be sensed by the stem cell. However, this may not describe the whole mechanism as signals decay with distance and may not reach the stem cell. Therefore, some other forms of communication between the stem cell and differentiated cells is necessary; we assume that such communication is important for maintaining tissue integrity. For this reason, we introduced entropy as a measure of global communication which also decays with distance but all differentiated cells communicate with the stem cell. Entropy thus is a property of the stem cell. A change in system entropy triggers the stem cell to initiate an autonomous damage identification and recovery process where it first starts to discern the general region of damage from the entropy change across the tissue regions (Fig 1b and c). Then, it triggers the local sensing sub-model, to activate the part of the AANN in the identified general damage region to find the exact location, size and shape of the damage. Finally, the stem cell migrates to the damaged location through the ECM to regenerate missing cells by asymmetric cell division (making a new differentiated cell while keeping its original form intact) until complete and correct recovery of the original tissue pattern.

##### 3.1.1.1 Global Sensing – damage region identification

This submodel senses the entropy change and scans the system by segments to find the general area of the damage (Fig. 1b and c). The difference between the current and original entropy informs the segment(s) that have received damage. Signalling entropy *E_i_* of the system defined as in Eq. (1) is a global measure of probability *p_ij_* or intensity of communication where the larger the distance *d_ij_* between the stem cell *i* and a differentiated cell *j*, lower the probability. *k* is the total number of differentiated cells.

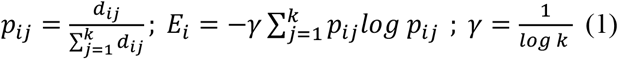

##### 3.1.1.2. Local Sensing – exact damage location identification

In local sensing, the stem cell initiates a local search for the exact location of damage in the already identified general area of damage by activating the corresponding region of the AANN to process information through local feedback connections to assess any missing neighbours. As illustrated in Fig. 2 and Eq. (2), each interior cell in the AANN has four neighbours that can communicate with it (Fig.2a) and the perceptron responds with 1 only if all neighbours are present and 0 otherwise (Fig.2b and c). The weights *w*_i_ indicate the strength of the incoming signals and we assume them to be equal in strength (*w*_i_ =1.0) as communication is only local and indicates simply the presence or absence of a neighbour. Therefore, the threshold for firing a perceptron is 4. Thus output of 1 means that all its neighbours are present (Fig.2c). In contrast, if the sum is less than the threshold, the output is zero indicating absence of one or more neighbours due to damage. After damage, interior neurons surrounding a damage have maximum of three inputs. As for the border of the original tissue, perceptrons have three inputs. In a damage case, border neurons near damage have maximum of two inputs. From these, the system can identify the damage border in the tissue interior (Fig. 2d) or at the boundary that can be used to recognise when to stop growing. These neighbour rules are properties of the tissue.

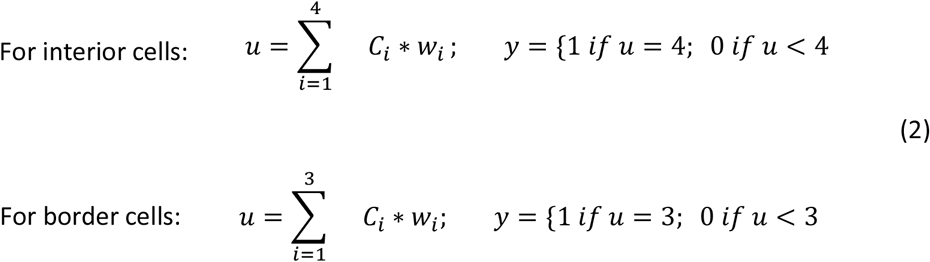

**Fig. 2.**
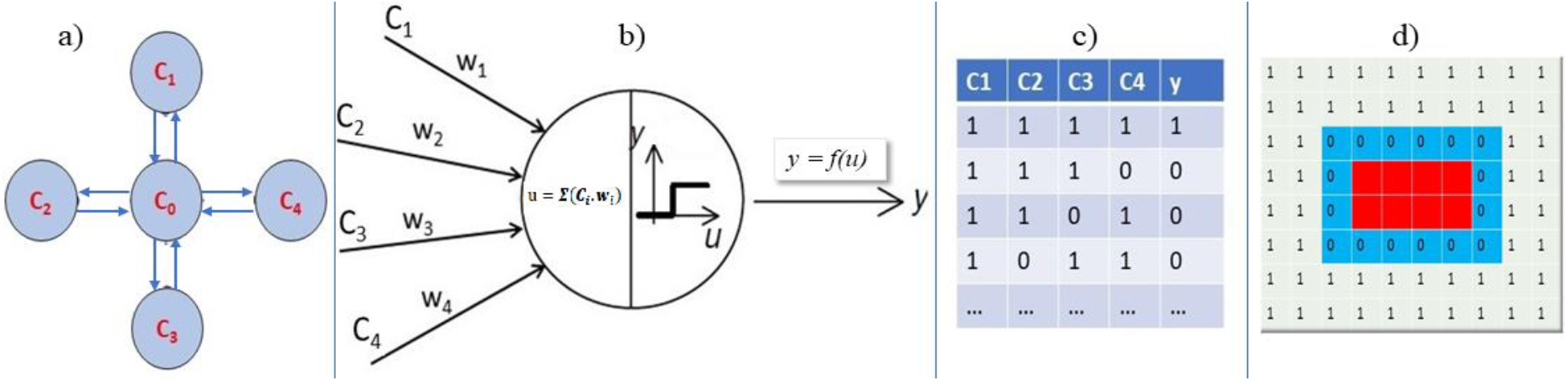
Perceptron communication and damage identification in the AANN – a) Two-way communication (local recurrent feedback) of a Perceptron with its neighbours in the AANN, b) Computation in a Perceptron, c) Sample data showing Perceptron response y to inputs Ci from neighbours, and d) Output of a damaged segment of AANN (red area) – 0 indicates a Perceptron sensing a missing neighbour and 1 indicates Perceptrons sensing the presence of the full set of neighbours.

##### 3.1.1.3 Regeneration

The single layer perceptron outputs help the system identify the boundary of the damage. In order to regenerate, the stem cell migrates to the damage location and fills out the damaged area by asymmetric cell division. In the case of a damage extending to the original border, it adds new cells until they align with existing border neurons. When a new cell is added next to a border neuron, the new cell becomes the marker for the next border cell and so on until the whole set of border cells is added and regeneration comes to completion with the renewal of the correct original form. Fig. 3 shows the operation of the tissue self-repair system for 4 types of damage (from small to large). Fig 3A shows the entropy change over the 24 tissue segments from the original state for the damage cases indicating that the stem cell senses very even small damages involving few cells (in fact, this system can recognise even single cell damages). This activates the local sensing with AANN that identifies the exact location of the damage (for example, red region in Fig. 3B (image 1) is the identified large damage in Fig. 3A(b)). The rest of the Fig. 3B illustrates the progression of recovery from this damage until complete and accurate regeneration. As stated earlier, this model can handle a large variety of damages; however, it requires that at least one string of cells from the stem cell to the border remains in the damaged tissue for recovering from extreme damages, such as losing a whole exterior region. To correct this, we introduce smart stem cells in the model extension in the next section.

**Fig. 3.**
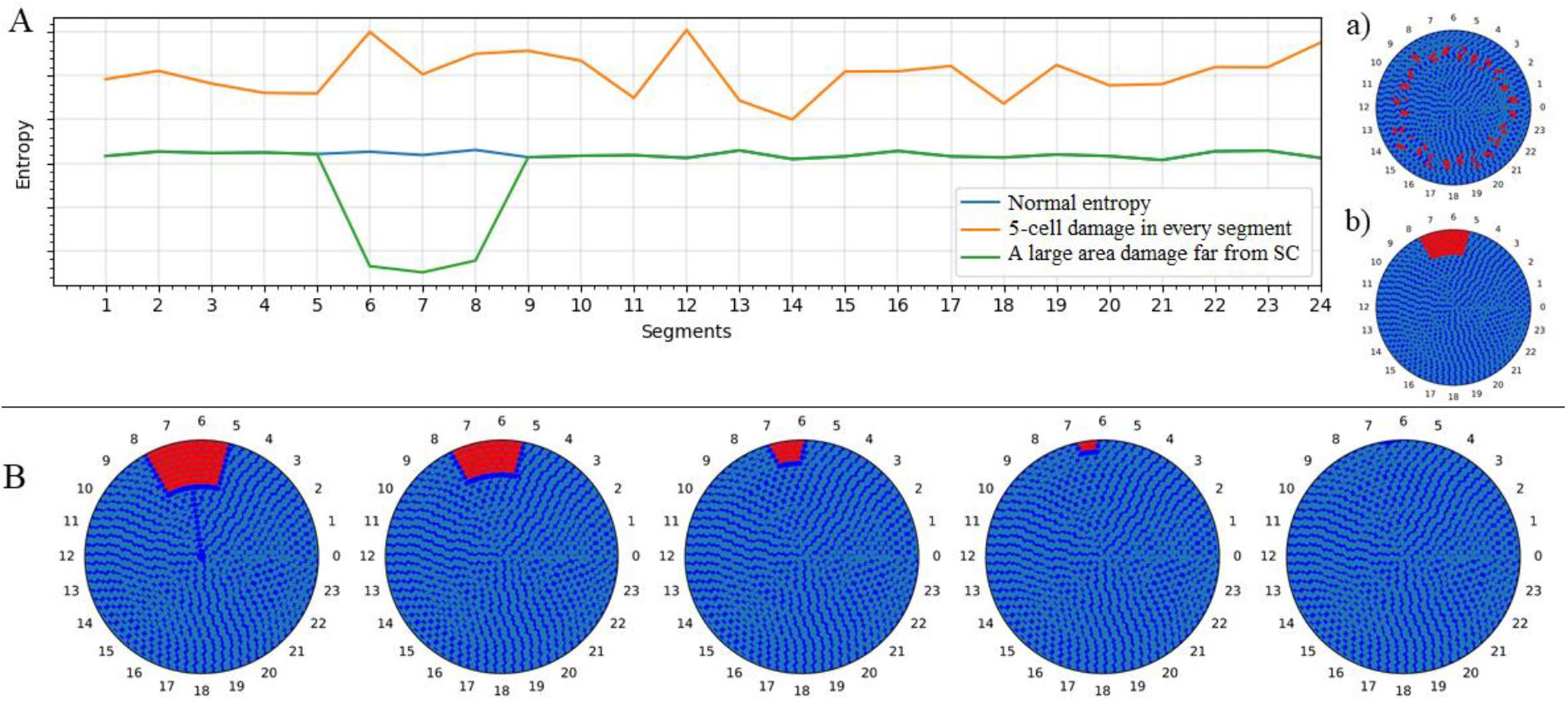
Entropy change and regeneration. (A) System entropy over the 24 segments of tissue for: no damage, and random deletions of 5 cells (as in (a)) in each segment, and 100 cells far from the stem cell (SC) (as in (b)); and (B) Progression of damage repair until correct completion (from left to right) for damage in Fig. 3A(b)

Also in the next section, the model is extended to more complex shapes such as triangular and rectangular shapes with smart stem cells and virtual fields and combine the shapes into an organism with a smart stem cell network and a collective virtual field to form the autonomous whole system regeneration model.

### 3.2 Extension of the base model: A Regeneration system with smart stem cells and information fields

We propose stem cells that remember the minimum information about their corresponding tissue patterns, that is stored in an information field surrounding the stem cells and shared as needed with other stem cells through the collective field. The information field is a mathematical abstraction over traditional components of tissues. We call them *intelligent* or *smart stem cells*.

#### 3.2.1. Circular tissue self-repair model with a smart stem cell

The circular tissue uses the same AANN structure and the tissue model shown in Fig. 1 but with a smart stem cell as in Fig. 4. The smart cell is basically the same as the previously described stem cell but with extra capabilities. Here, as the stem cell communicates with the differentiated cells, it uses this information to recognise the boundary of the tissue pattern and extract minimum pattern information required for regeneration. Specifically, the smart stem cell keeps radius *d* in the information field, that allows it to regenerate the pattern from any damage, thus releasing it from the previous constraint of having to retain minimum information in the damaged tissue. It is important to note that in most cases of damage, system would regenerate without tapping into the field. Stored information will be used only in extreme cases where the whole external region goes missing. This is not unlike the situation where, in most cases of damage to a building, a structural engineer would repair it by taking cues from the undamaged part of the structure; however, when the damage is severe to the extent that it creates open-ended boundaries, they refer to the original plan to repair the damage.

**Fig. 4.**
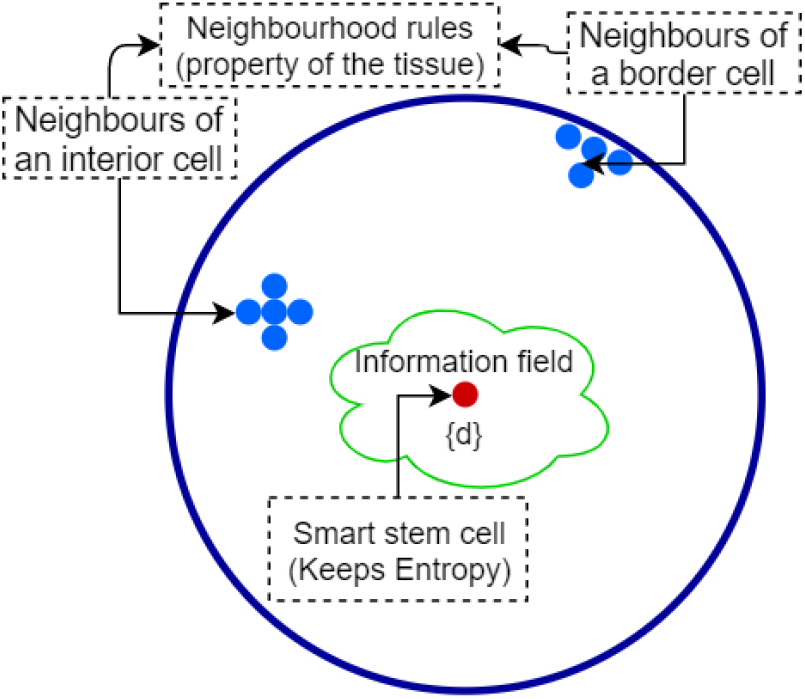
A circular tissue with a smart stem cell and surrounding information field

#### 3.2.2 Triangular tissue and Self-repair model

We develop a self-repair system similar to the one in Figure 1c for the triangular tissue but different neighbourhood rules apply to the AANN and the smart stem cell (Fig. 5). The neighbourhood rules are: the number of neighbours of interior cells, border cells and corner cells are four, three and two respectively. From these rules, perceptrons in the AANN can collectively establish the damage borders. Tissue is divided into four segments for monitoring its state through entropy (Fig. 7a). Without a smart stem cell, the system can regenerate from many damages except for the extreme cases where one whole side or whole exterior region receive damage creating open ended boundaries. To cover for these cases, we introduce a smart stem that carries some minimum shape information in the information field, extracted from its communication with the differentiated cells: the length of the sides (d), aspect ratio of sides (AR=1), the number of corners (n=3). (Entropy is a property of the stem cell and neighbour rules are properties of the tissue). With this information, it achieves complete regeneration with the help of the AANN. In the next section, we demonstrate the operation of this tissue repair system.

**Fig. 5.**
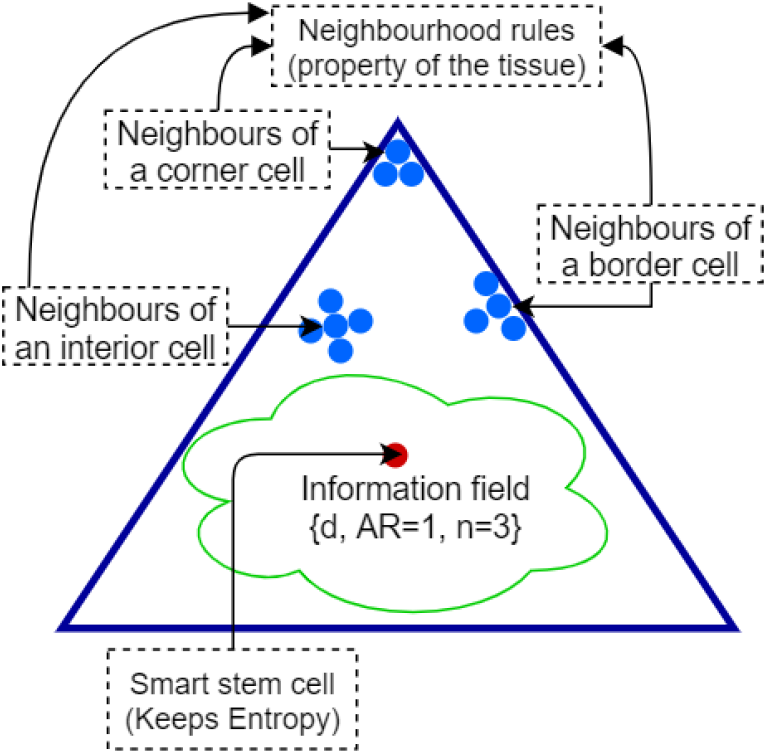
A triangular tissue with a smart stem cell and surrounding information field

##### 3.2.2.1 Two cases of regeneration of triangular tissue

We demonstrate how the system recovers from two types of damages shown in Fig. 6. In the first case, a part of the tissue is missing (Fig. 6a), and in the other, the tissue incurs severe damage leaving a small fragment with the stem cell (Fig. 6b).

**Fig. 6.**
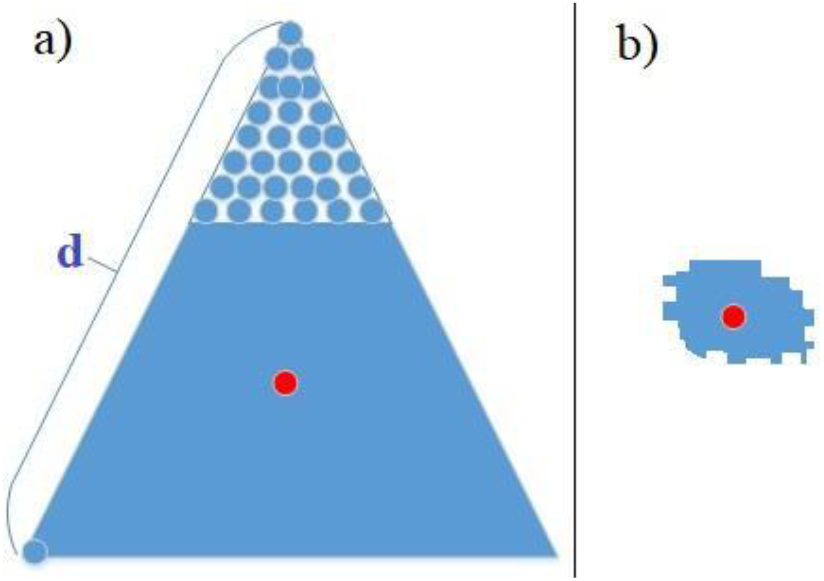
A triangular tissue with two cases of damage. (a) tissue loses a part of its structure, and (b) tissue incurs severe damage

###### a) Tissue loses a part of its structure

For the first damage case, Fig. 7 shows the entropy change and Fig. 8 shows the repair process. Global Sensing based on entropy identifies the damage regions of 3 and 4 (Fig. 7b) from the entropy of the four segments shown in Fig. 7a, and from this, AANN identifies the exact damage location and the border cells of the damaged area (Fig. 8c) using rules for interior and border cells shown in Fig. 5. Then, the stem cell moves to this location and produces new cells row by row along the damaged border to regenerate the pattern without tapping into the field as it can use the existing pattern information in the damaged tissue. Specifically, new border cells are added that are guided by current border cells of the structure and neighbourhood rules for the border. When a border cell aligns with two border edges and it does not satisfy the border rules, then this border cell becomes a corner cell that stops regeneration (Fig. 8d).

**Fig. 7.**
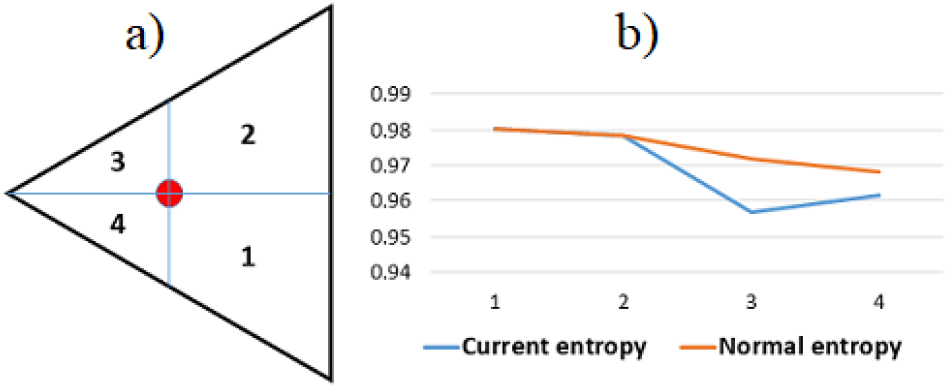
Global sensing of the self-repair model of triangular tissue. (a) Triangular tissue with stem cell and tissue segments used for entropy calculation, and (b) Original and current entropy across the segments for the damage case in Fig. 6(a)

**Fig. 8.**
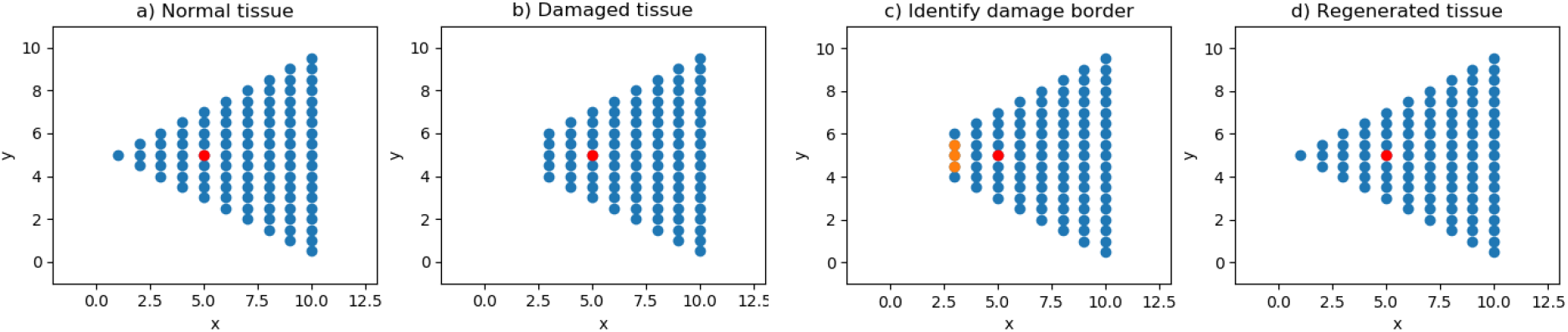
Regeneration steps for damage in Fig. 6(a)

###### b) Tissue incurs severe damage leaving a small part with the stem cell

Fig. 9 illustrates the progression of regeneration to restore the original form from the severe damage in Fig. 6b. Damage identification is the same as before with Global Sensing and Local Sensing. The first image in the second row of Fig. 9 shows that all 4 segments have received damage. As this is an extreme damage to tissue exterior, the system taps into the information field. The stem cell initially produces new cells to form a new small triangular tissue (Fig. 9b) that then incrementally grows until completion (Fig. 9c-d) according to the above neighbour rules and minimum geometry information in the virtual field. The rest of the figures in the second row shows the restoration of tissue entropy to original state during regeneration.

**Fig. 9.**
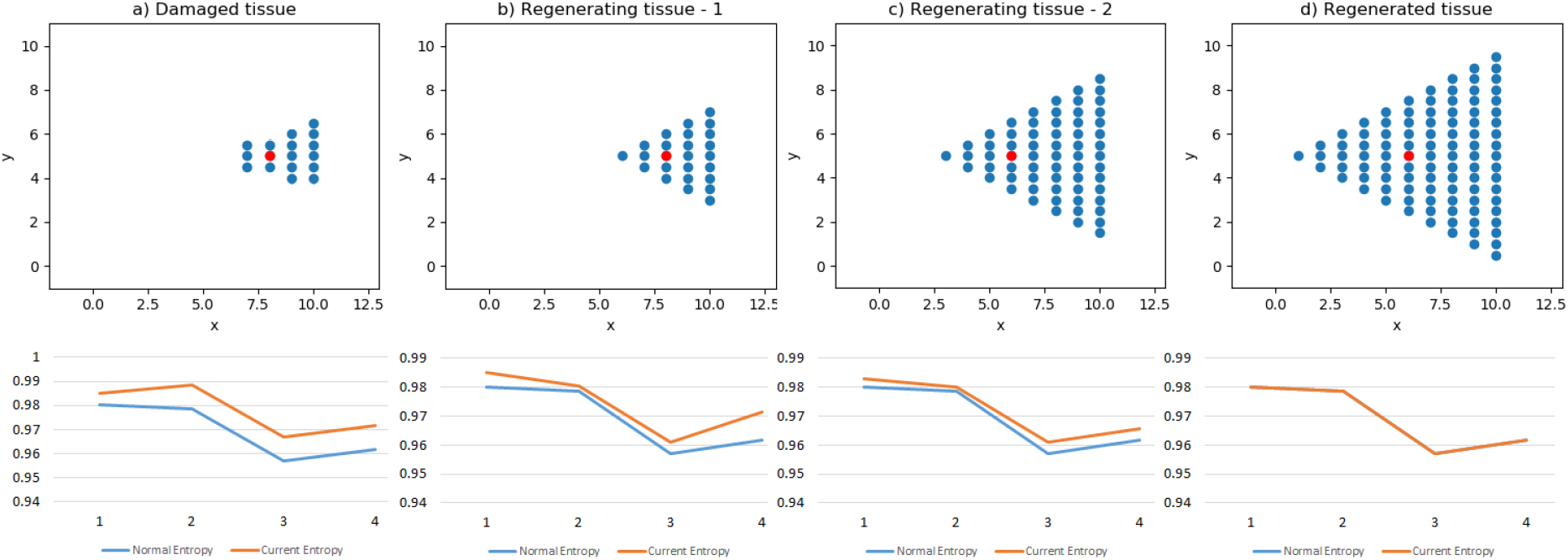
Regeneration of a triangular tissue from a small remaining fragment with intact smart stem cell. First row shows stages of regeneration of the pattern until complete recovery, and the second row shows corresponding tissue entropy

#### 3.2.3 Rectangular tissue and Self-repair model

We develop the self-repair network for a rectangular tissue using the same method but different neighbourhood rules apply to the AANN and smart stem cell. Neighbourhood rules are: the number of neighbours of interior cells and border cells is four and three, respectively (Fig. 10). Corner cells have three neighbours where two of them are border cells that are also neighbours of the third cell. Smart stem cell carries some minimum shape information in the information field: width (d), aspect ratio of width to height (AR=3), the number of corners (n=4). In this tissue, the system can recover from any damage without tapping into the field as long as a whole side or whole exterior does not receive damage. For such damage to exterior, it taps into the field for pattern information.

**Fig. 10.**
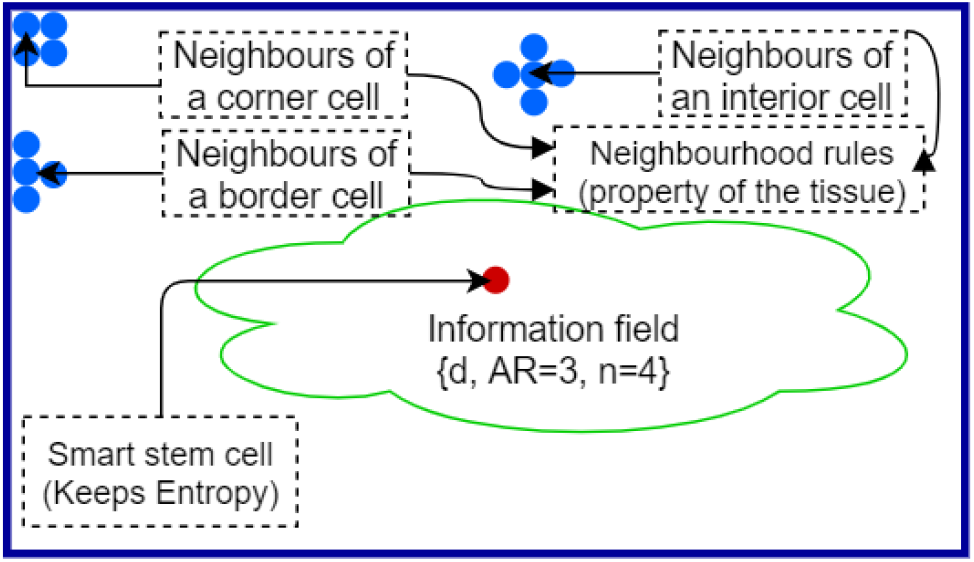
A rectangular tissue with a smart stem cell.

##### 3.2.3.1 Two cases of regeneration of rectangular tissue

We apply the tissue repair model to 2 cases of damage; a damage where the tissue loses one side (Fig. 11a), and a severe damage where the whole exterior region is lost leaving only a tiny fragment with the stem cell (Fig. 11b).

**Fig. 11.**
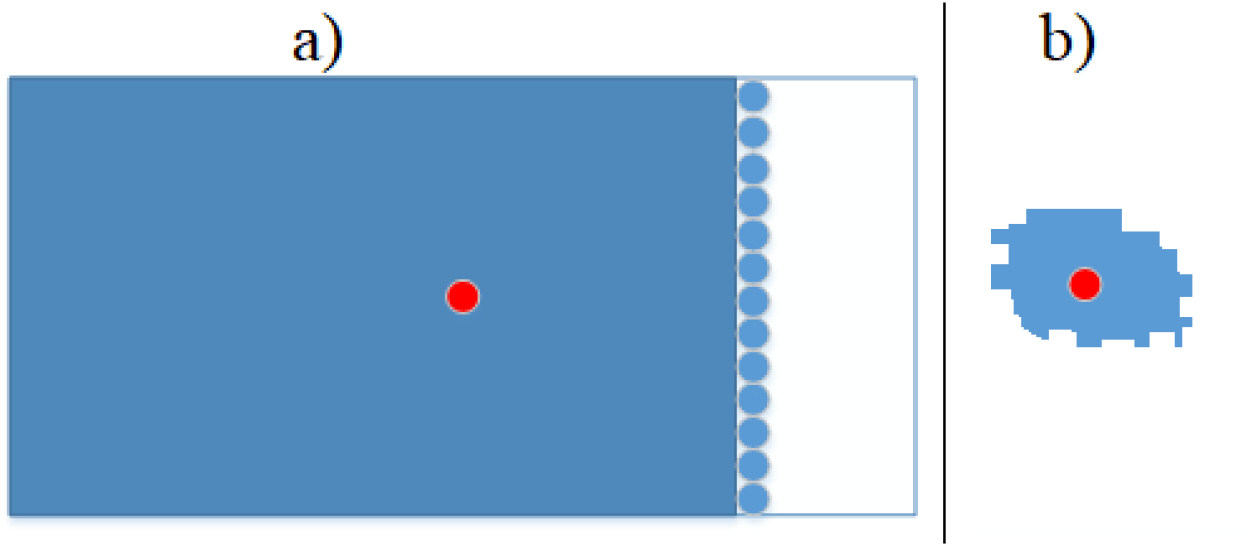
Two cases of damage – (a) damage to one side and (b) severe damage leaving a small fragment with the stem cell

As in the triangular case, system identifies the damage region with Global Sensing using entropy and finds the location and size of damage through Local Sensing of missing neighbours in the AANN and regenerates completely with the above neighbour rules and pattern information. As a side has received damage, the system taps into the field for required pattern information. Fig. 12 shows the damaged tissue, segmented tissue and entropy before and after damage as well as AANN detected damage border and the regenerated tissue for the first damage case.

**Fig. 12.**
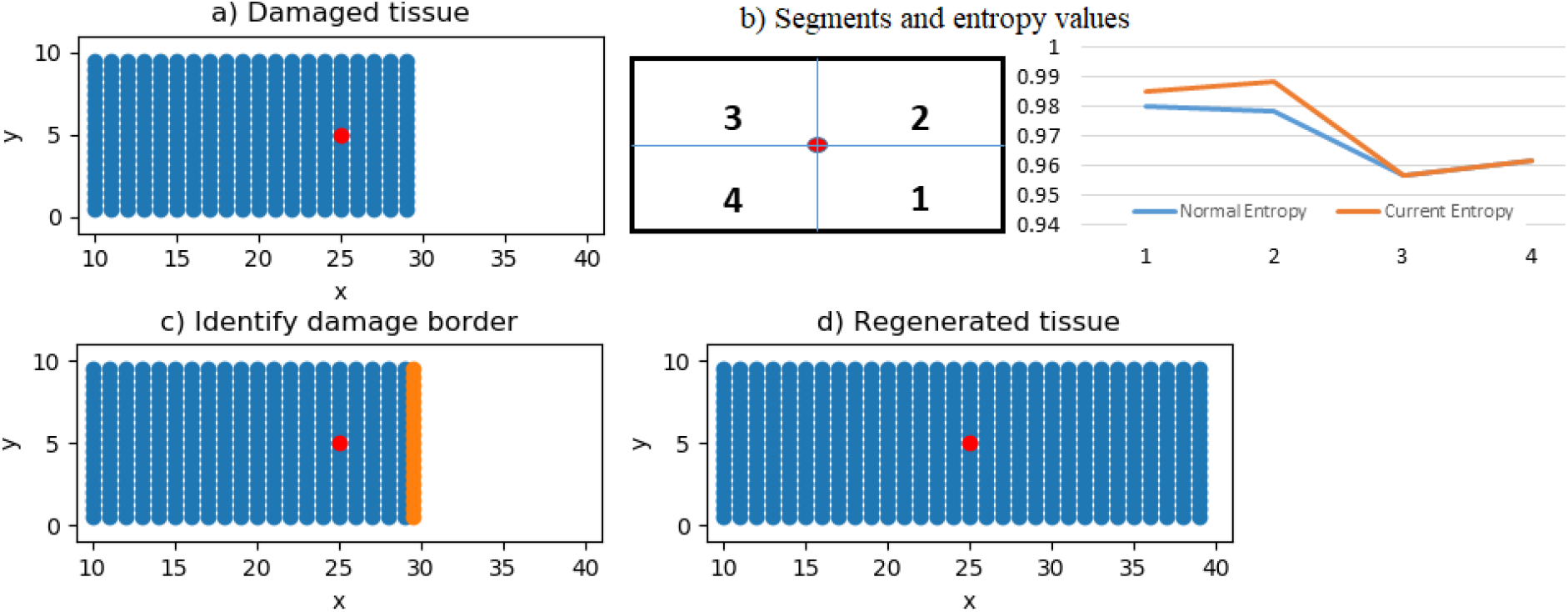
Entropy change, AANN identified damage border and regenerated tissue for the damage in Fig 11a.

Fig. 13 shows the stages of progression of regeneration to complete recovery for the second damage case in Fig 11b. As the whole exterior has received damage, pattern information is accessed from the field.

**Fig. 13.**
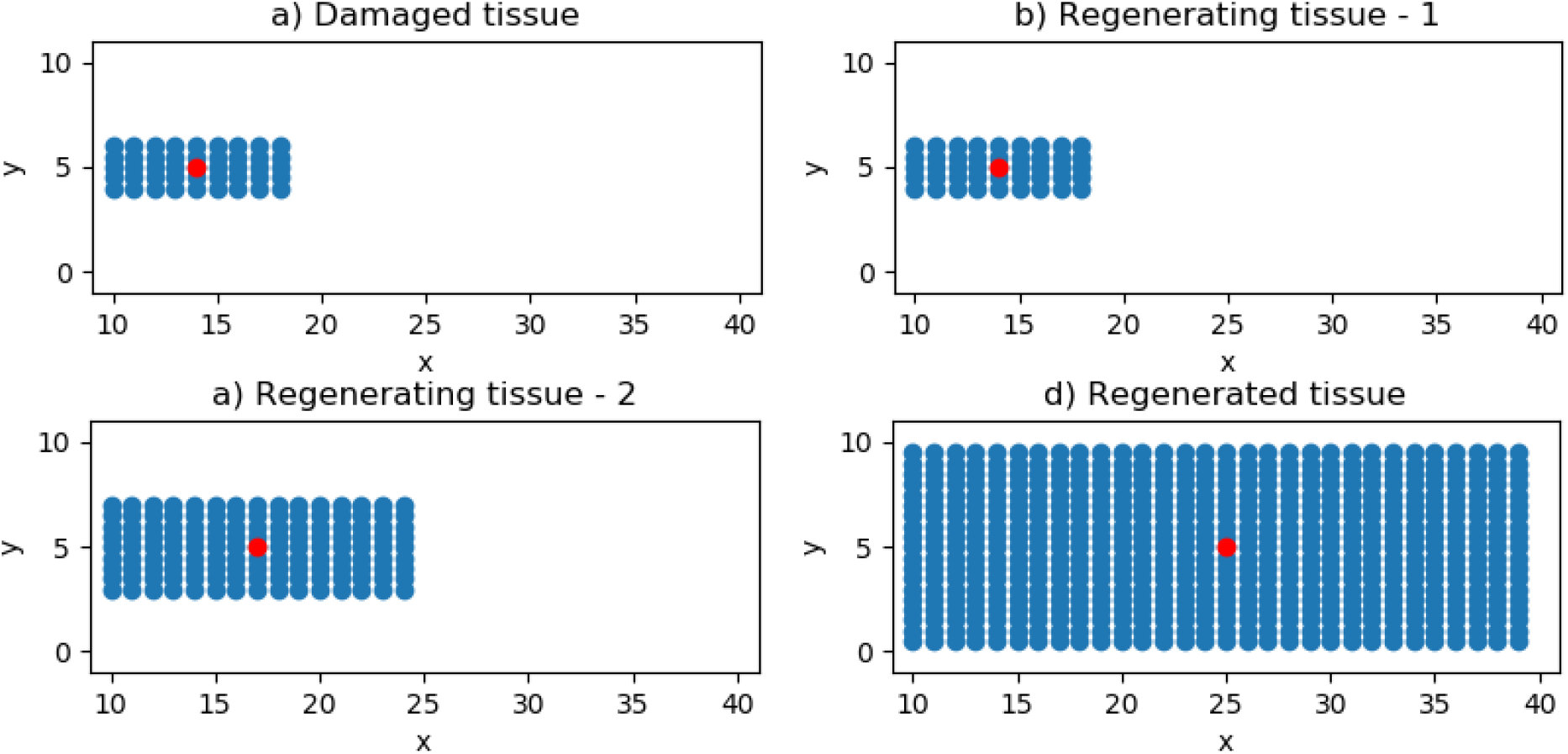
Regeneration of a rectangular tissue from a small fragment as in Fig. 11b

## 4. Whole system regeneration model

### 4.1. Creating a virtual organism from individual tissues with overlapping information fields of stem cells

The individual tissues in the previous section maintain the tissue integrity. When damage happens in the system, the smart stem cell with the cell collective detects and repairs the damage immediately. We now create a virtual organism by combining these individual parts with smart stem cells to represent a system like planarian. Specifically, we assemble triangular and rectangular tissues into a simple organism shown in Figure 14a with head, body and tail. It shows the shared information field around the organism (with information presented in vector form) as applicable to the three tissues. The shared field enables regeneration from any damage anywhere in the system as long as one (any) stem cell remains.

**Fig. 14.**
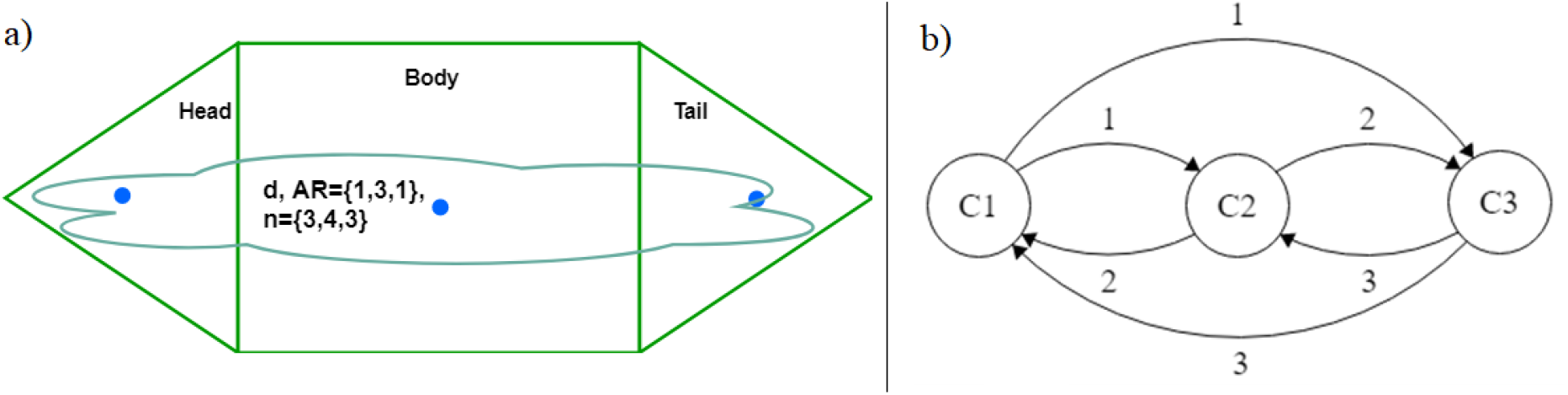
Two levels of the whole system regeneration model. a) Form of the virtual organism with head, body and tail tissues and shared collective information field, and b) Stem cell network of head (C1) body (C2) and tail (C3). AR={1,3,1} (aspect ratio of {head, body, tail}) is width to height ratio for rectangular tissue and ratio of sides for triangular tissue; d is the length of sides of triangular tissue and height of the rectangular tissue; n={3,4,3}is the number of corners of {head, body, tail}.

### 4.2 Overall model framework

The overall framework and the operational flowchart of the whole system regeneration model are shown in Fig. 15 and 16, respectively. It achieves complete regeneration through two levels of operation:

**Fig. 15.**
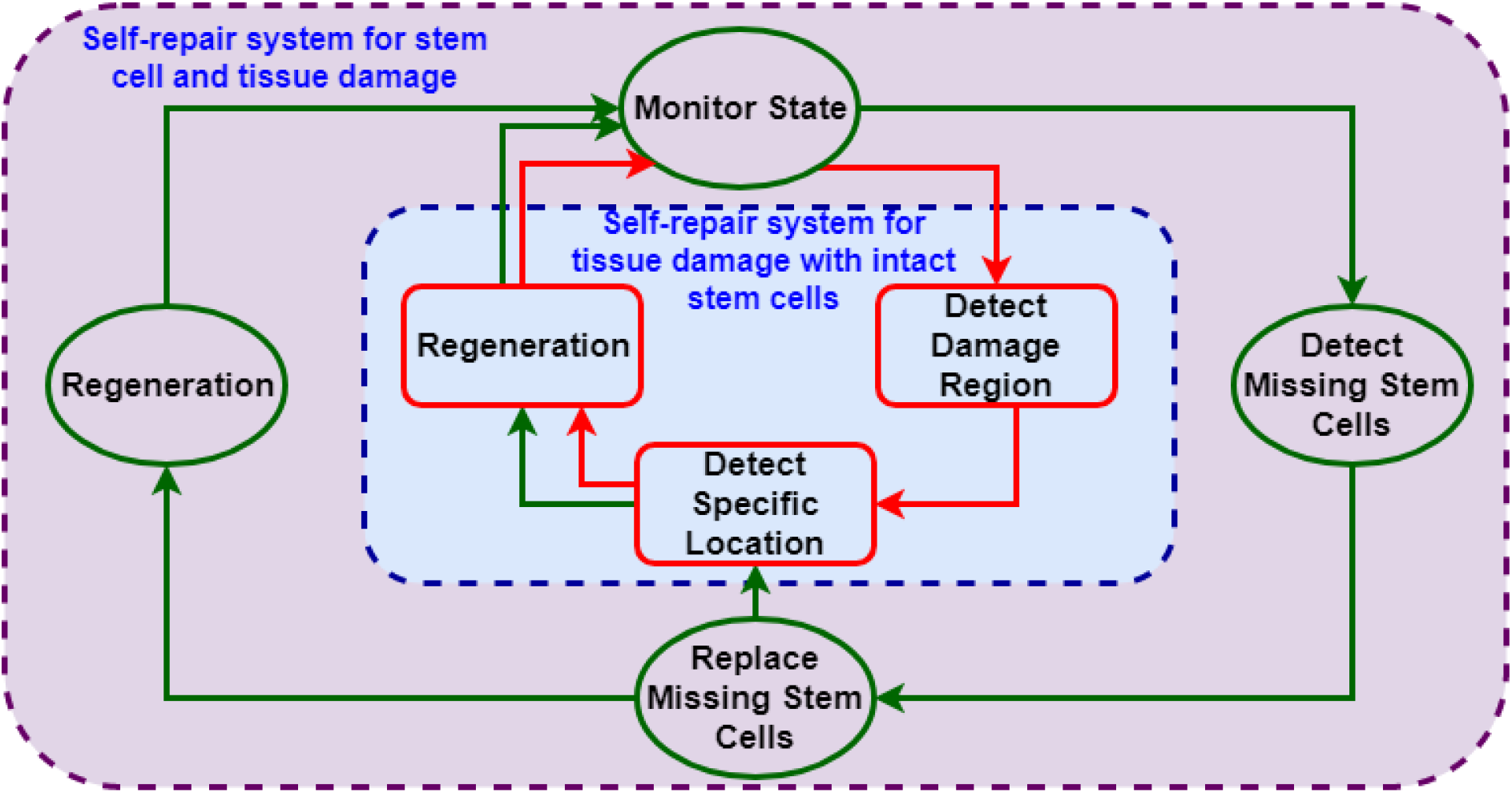
The System Framework for full organism regeneration after tissue and/or stem cell damage

**Fig. 16.**
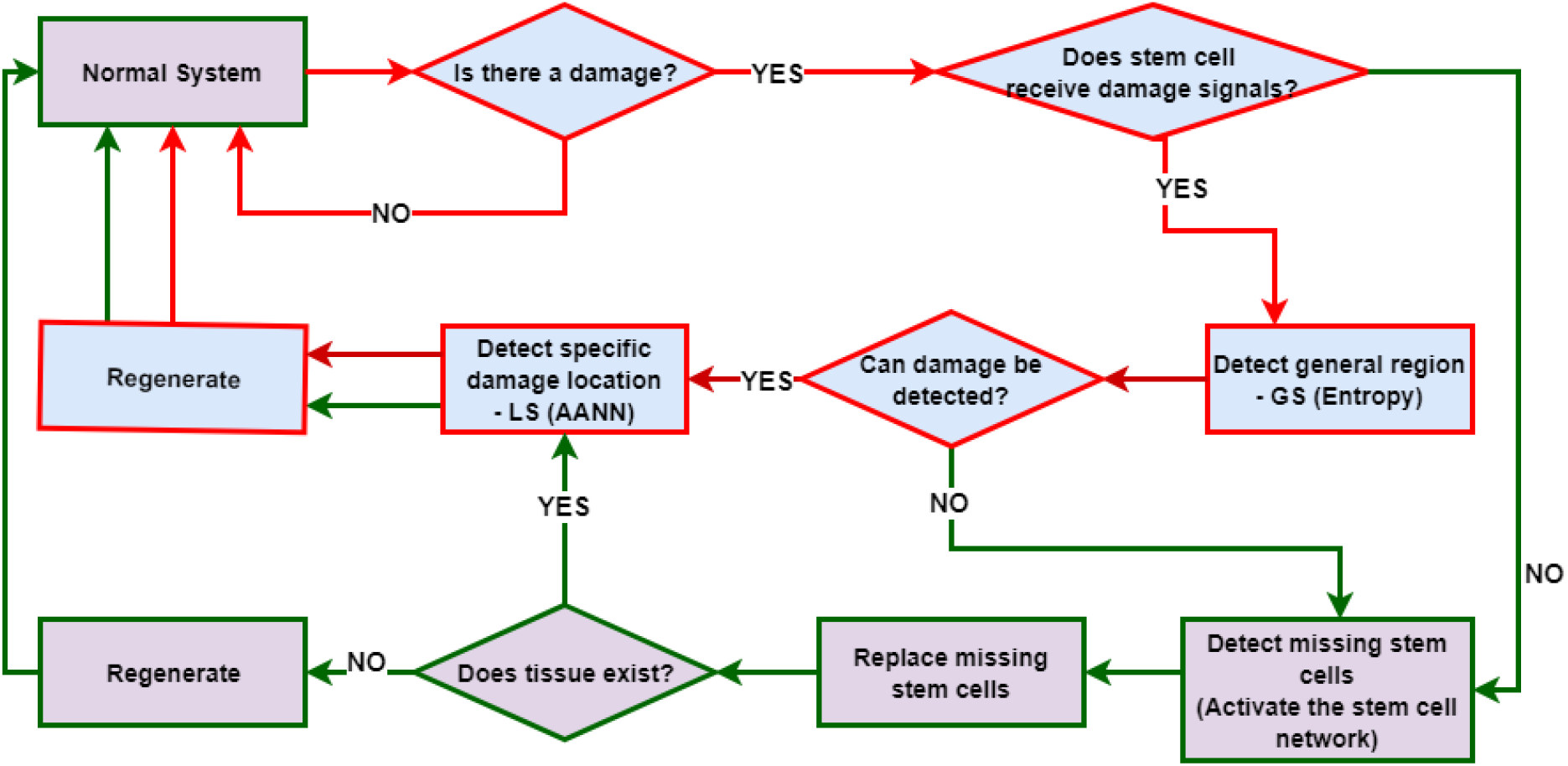
The flowchart of the whole system regeneration model

**Level 1:** This sub-model represents individual tissue self-repair systems for cases where stem cells are intact. It consists of individual tissues with smart stem cells (Fig 14a) and AANN, with Global and Local Sensing features (inner blue box in Fig. 15 and blue items in Fig. 16).

**Level 2:** This is the stem cell repair system for recovering from the loss of stem cells along with *whole* tissues (head, body or tail) (sub-model 2) (outer purple box in Fig. 15 and purple items in Fig 16). It consists of a stem cell network with overlapping information fields (**Fig. 14a and b**). Here, stem cells communicate with each other through signals weighted according to their location identifier (1, 2, or 3). We describe in detail the development of the stem cell repair network model in the next section.

The idea of the new framework is that when there is only tissue damage leaving stem cells intact, only Level 1 of the model is activated where the stem cell senses the damage through entropy change, detects the general damage region and triggers the AANN to find the exact damage location. Then the stem cell migrates to the damage region and repair it with the help of the AANN as shown previously for individual tissues (red arrows in Fig. 15 and 16). This system taps into the information field only for extreme tissue damage.

For any damage involving stem cells and whole tissues (head, body etc.), Level 2 is activated where stem cells communicate with each other through the stem cell network and senses and regenerate missing stem cells using information in the shared (overlapping) fields (green arrows in Fig 15 and 16). The shared information (e.g., minimum pattern information) can be accessed by all smart stem cells. This idea is used in the production of new stem cells. Specifically, a lost smart stem cell is replaced with a new smart stem cell with similar characteristics as before by the neighbour smart stem cell that transfers the shape information from the shared field into the new smart stem cell. The new stem cell then carries out complete regeneration of the damaged tissue using the received pattern information, and a new tissue AANN is re-established along with neighbour rules. The stem cell then restores tissue entropy.

When more than one stem cell is lost, new smart stem cells are produced by the remaining stem cell using the pattern information in the field for concurrent and seamless regeneration of the missing parts and whole organism. The two levels collaborate when only a part of the tissue (not the whole tissue) is lost with stem cells, as shown by the green arrows flowing through both submodels in Figure 15 and 16 where after replacing the missing stem cell(s), the corresponding tissue model(s) is activated with entropy based global sensing and AANN based local sensing to detect damage and its border and execute full regeneration of the damaged tissue.

### 4.3 Operation of the whole system regeneration model: Normal system operation and damage detection

In the intact system, the stem cell network and individual tissue models work normally. Tissue models are activated for damages with intact smart stem cells and stem cell network is activated for tissue damages with loss of stem cell(s). We explore three potential approaches for stem cell repair network using Automata, Neural Networks or Decision Trees as illustrated below. The reason for three approaches is to test them in order to find the most efficient and robust method.

#### Automata

Automata is a system of nodes that communicate according to string grammar rules (Table 1). Each node (stem cell) sends signals (1 or 0) to other nodes as shown in Fig. 14b. Signals are weighted based on a cell identifier (1, 2 or 3).

**Table 1.** String grammar rules

|  |  |
| --- | --- |
| 1. C1 -> 1 C2 | 2. C1 -> 1 C3 |
| 3. C2 -> 2 C1 | 4. C2 -> 2 C3 |
| 5. C3 -> 3 C1 | 6. C3 -> 3 C2 |

##### • Normal operation

Normal system operation is recognized when all stem cells are present as defined by the string grammar rules shown in Table 1 for sending and receiving information. Basically, rules describe how cells communicate with each other and weights identify the cell where the communication originates. In this system, relative location based cell identifier (1, 2 or 3) is used as weights.

##### • Damage detection

If the damage involves missing stem cell(s), string grammar rules fail and missing stem cells are identified.

#### Neural networks

##### • Normal operation

Each stem cell is a linear neuron that receives signals from the other stem cells as shown in Fig.17. System operation is normal when the output of each stem cell records a maximum value as explained in Equations (3, 4, 5) and Fig. 18.

**Fig. 17.**
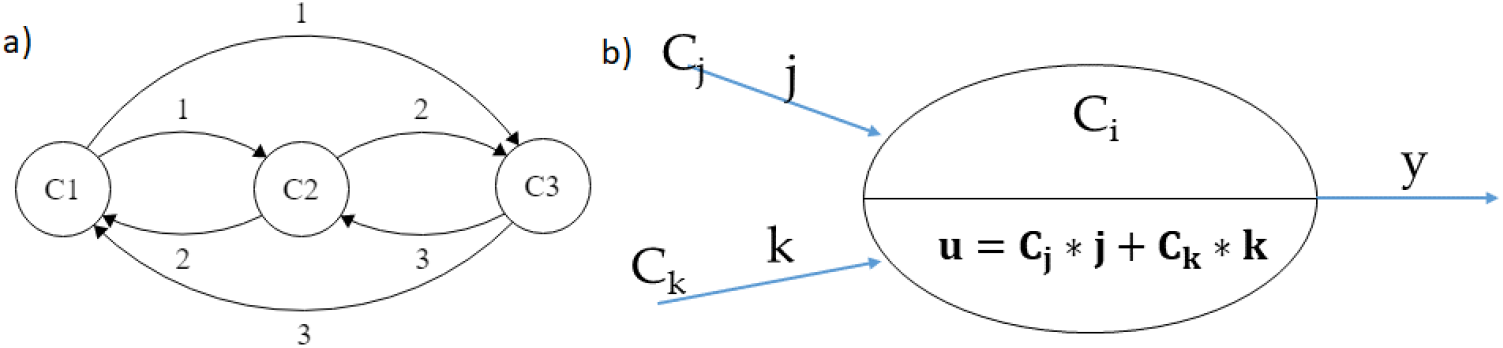
a) Network of stem cells and b) Neural network model of a stem cell

**Fig. 18.**
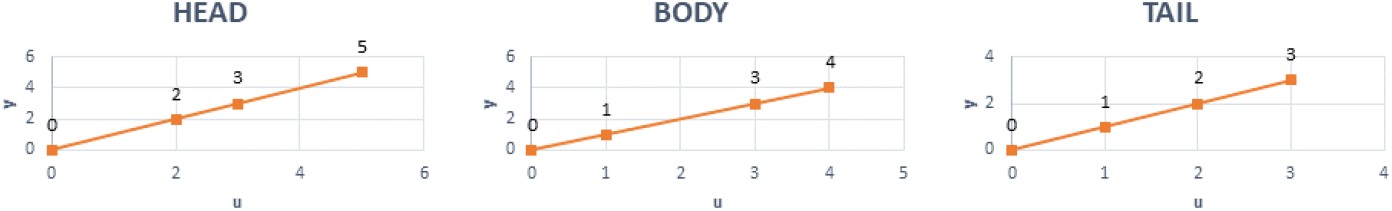
Neural network output of individual stem cells

Output of each stem cell neuron is as follows:

<u>Head stem cell</u>:

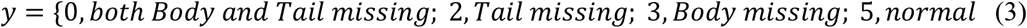

<u>Body stem cell</u>:

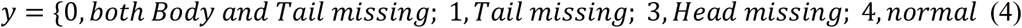

Tail stem cell:

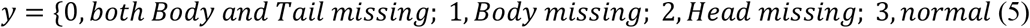

##### • Damage detection

When damage happens resulting in missing stem cells, the neuron output is smaller than that of the normal system output according Eq. 3–5 and Fig. 18. This way, missing stem cells are identified.

#### Decision tree (using J48 algorithm in Weka)

In computer science, a definition of a decision tree is “a flowchart like tree structure, where each internal node denotes a test on an attribute, each branch represents an outcome of the test, and each leaf node holds a class label. The paths from root to leaf represent classification rules.” (Han, Kamber, & Pei, 2006). A decision tree is a model that helps determine the state of a system depending on decision rules. In our model for example, each stem cell receives two signals from the other stem cells. A decision tree model is built to classify the states of the system. Fig. 19b shows the decision tree for the stem cell network in Fig. 19a, where each leaf is an output of the model stating whether the system is normal or some parts are missing depending on the inputs received by each stem cell as shown in Fig. 19c where j and k values are as in Fig. 19a.

**Fig. 19.**
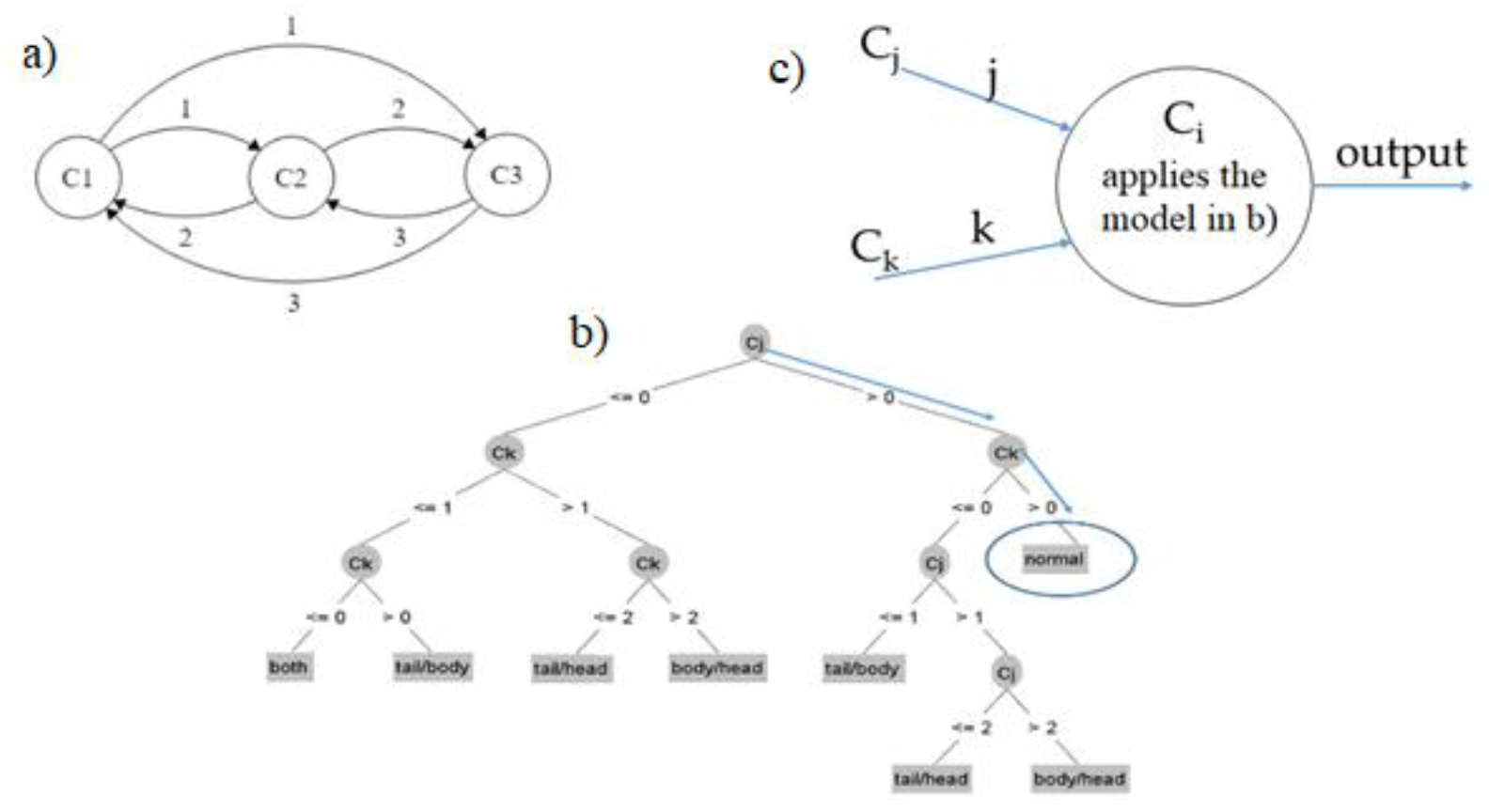
Decision tree model for damage detection. a) Stem cell network; b) Decision Tree model and c) Implementation of the model for each stem cell.

##### • Normal operation

System operation is normal when all stem cells are present returning an output corresponding to ‘normal’ as indicated by the direction of arrows in Fig. 19.

##### • Damage detection

When a damage happens to stem cells, the model returns an output indicating the damaged part (head, body or tail).

For example, head stem cell applies the decision tree model with two input signals from body stem cell and tail stem cell. There are three outcomes here:

— **Normal system:** C_j_=2, C_k_=3; For this case, the system follows arrows (1) in **Fig. 20**, and reaches the output for the **normal** case.
— **Tail is missing:** C_j_=2, C_k_=0; the system follows arrows (2) in **Fig. 20**, and reaches the output **tail/head** stem cell damage meaning that either **tail or head stem cell** is damaged. Because head stem cell is working in this case, tail stem cell is missing.
— **Body is missing:** C_j_=0, C_k_=3; Following arrows (3) in **Fig. 20**, the system reaches the output **body/head** stem cell damage. Because head stem cell is working in this case, body stem cell is missing.

**Fig. 20.**
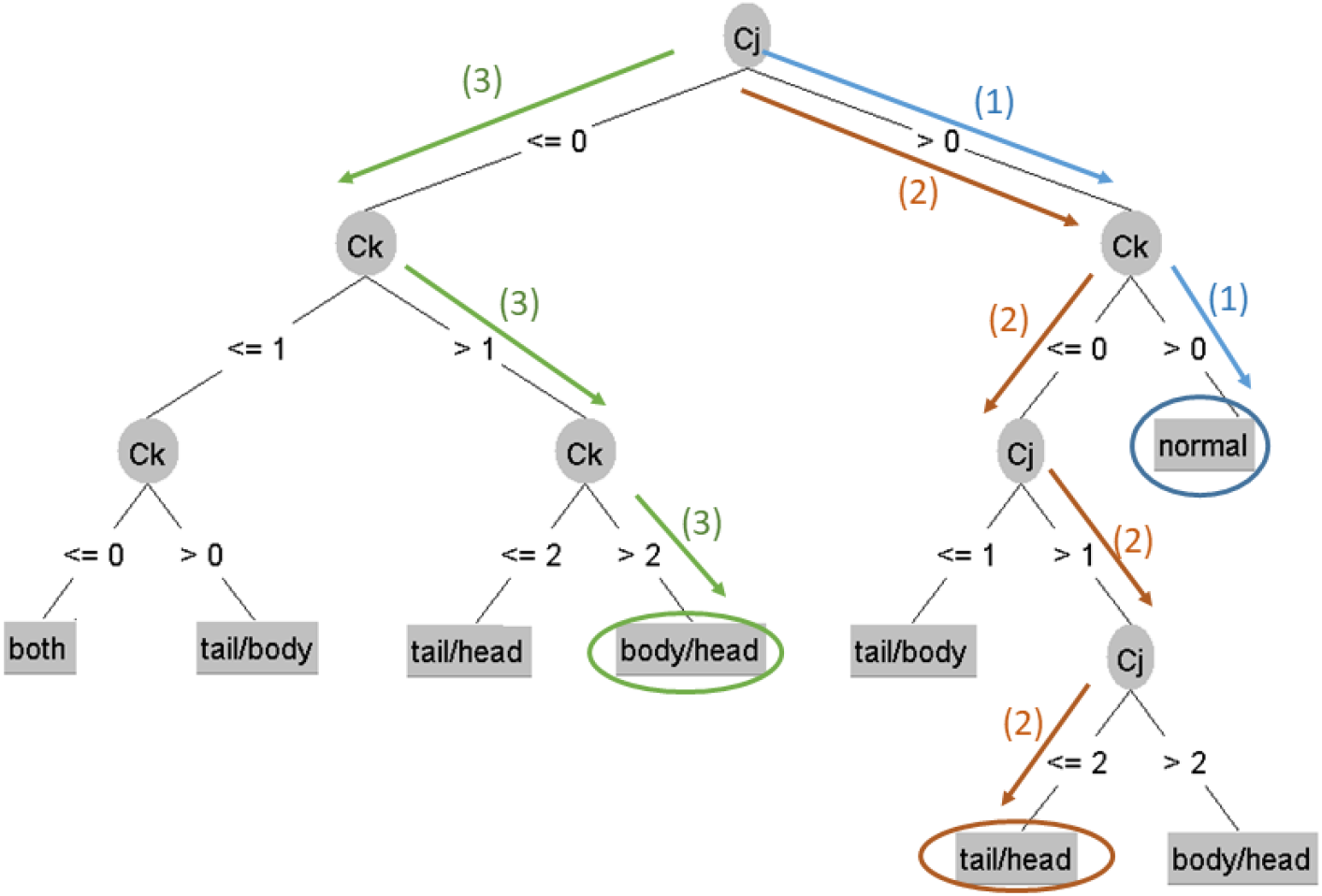
Decision Tree model for stem cell network for normal system and damage detection

## 5. Implementation of the new regeneration framework: Examples of regeneration

In this section, we demonstrate the ability of the whole system regeneration model to recover from any damage. We consider two main types of damage: partial tissue damage with intact stem cells; and severe damage involving loss of both tissue and stem cells anywhere in the organism.

### 5.1. Only tissue damage with intact stem cells: system regenerates completely

This type of damage is simple in that only a part of the organism is missing but smart stem cells are intact. It means that a group of differentiated cells is cut off while the smart stem cell of this part remains as in **Fig. 21** where the virtual worm is depicted in a plane with a set of coordinates (x, y). The red and blue dots are smart stem cells and differentiated cells, respectively.

**Fig. 21.**
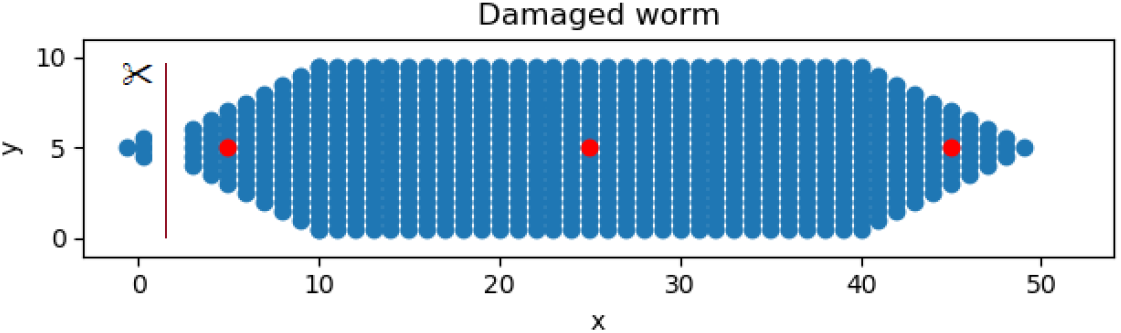
Virtual worm with a missing fragment of head

This case does not require accessing pattern information from the field and regeneration proceeds as follows. Damaged cells cease sending signals to the nearest smart stem cell that senses the resulting entropy change and activates the sub-model for tissue repair (the blue part with red arrows in **Fig. 15** – see also the part with red arrows in the flowchart in **Fig 16**) to initiate the following steps: (i) calculate entropy change, (ii) activate AANN, (iii) identify borders and (iv) repair damage. The small part of the tissue that was separated from the head tissue does not contain a smart stem cell so it cannot regenerate. The other remaining part with smart stem cells does regenerate the head fully and accurately. The first step is that the smart stem cell compares the stored and new value of entropy of each segment to identify general damage region. The graph in Fig. 22b) shows that the damage occurred in two segments 3 and 4 (i.e., at the front of the head). Then, the smart stem cell activates the part of the AANN of the head tissue corresponding to these segments to identify the border of the damage (Fig. 22c) using perceptron rules. The last step is that the smart stem cell migrates to the damaged border and regenerates missing cells using rules for border. The regeneration proceeds until the damaged part is regenerated returning the worm to the original form as shown in Fig. 22d.

**Fig. 22.**
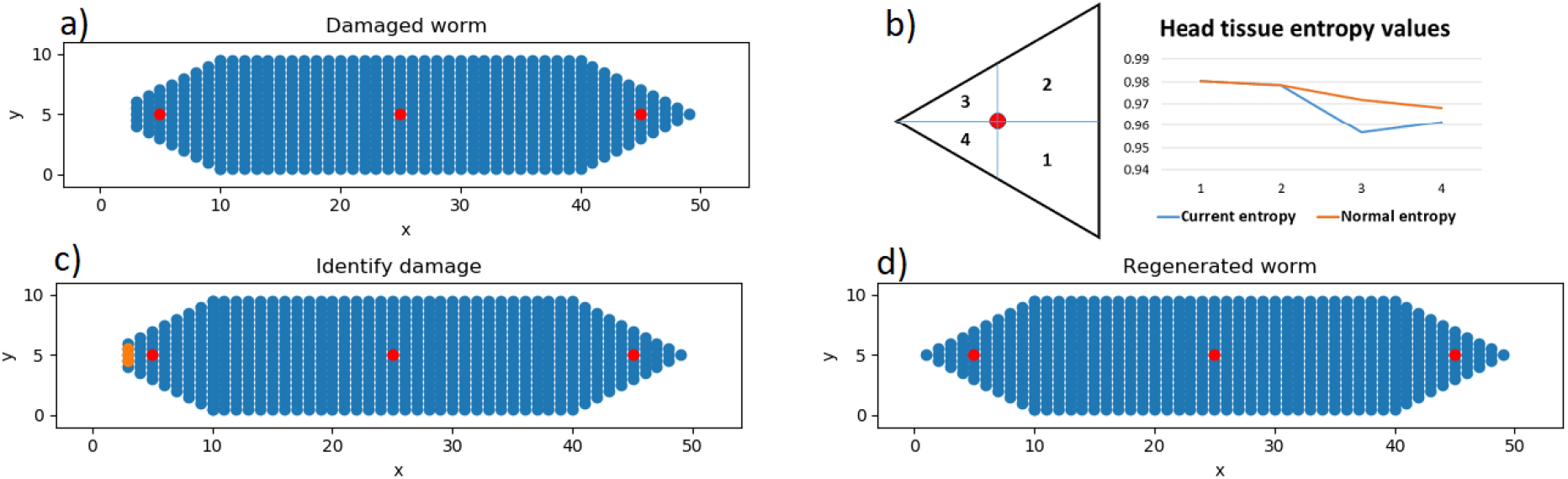
Head regeneration process: a) Damaged worm, b) Segmented tissue and entropy, c) Identification of damage border, and d) Regenerated worm. (x and y are coordinates)

### 5.2 Organism receives both tissue and stem cell damage

The main idea for this case is that the smart stem cells can be amputated with the surrounding tissue. We consider two damage situations for this case with an organism fragmenting into two parts: (i) along a tissue boundary; and (ii) encompassing tissue and stem cell damage in the body interior.

#### 5.2.1 Organism fragments into two parts along a tissue boundary

Here we consider a damage along the head-body boundary that fragments the organism into two parts with intact stem cells as shown in Fig. 23. The main question is: how does each part regenerate into a whole organism as the original one, resulting in two identical worms? We explain the process of regeneration below.

**Fig. 23.**
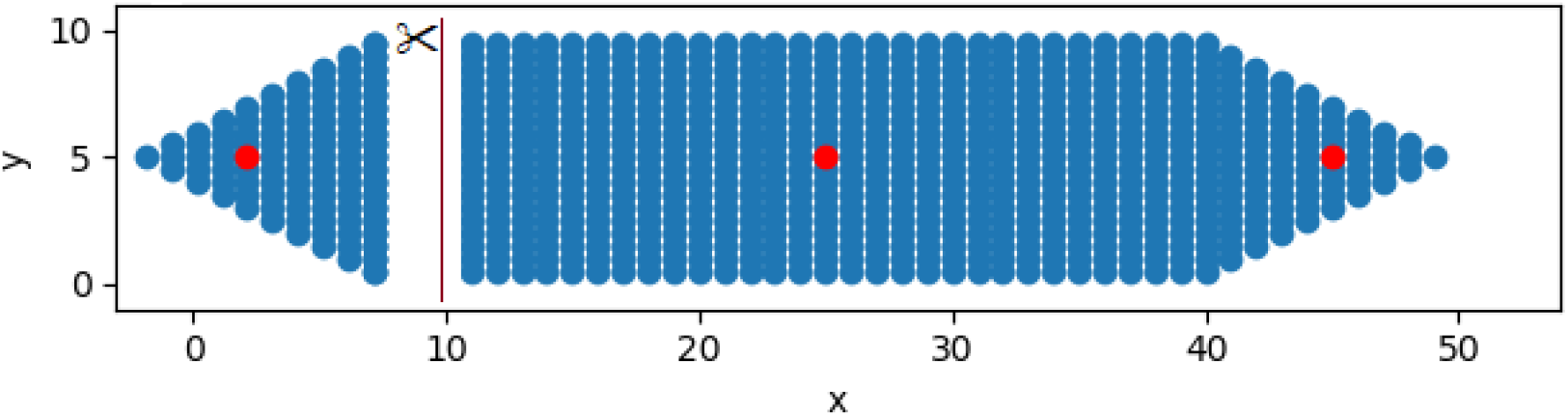
Virtual worm damaged along the head-body boundary with intact stem cells.

As the damage has left an open ended boundary, the system taps into the field for minimum pattern information. Both parts regenerate by activating the repair system as follows (follow the green arrows in Fig. 15 and 16): After amputation, smart stem cells in both parts check for any local damage as described in the case above for tissue damage. If there is no local damage, as is the case in this example, it will activate the stem cell repair model (purple part with green arrows) in Fig. 15 (and green arrows and items in Fig. 16) and follow three steps: (i) identify missing smart stem cell(s), (ii) replace missing smart stem cell(s), and (iii) regenerate missing tissues concurrently and re-establish AANN. We explain the regeneration of each missing part below.

##### 5.2.1.1 Head regenerates body and tail

###### Identify missing stem cells

From the operation of the stem cell network, head should recognize that the body and tail stem cells are missing in the organism. Here, we consider the three approaches selected for stem cell network to detect and identify missing smart stem cells. First, using Automata: head stem cell receives no signals from other smart stem cells. Then it knows they are missing based on the string grammar rules in **Table 1**. Second, using Neural Networks: as head stem cell receives no signals from others, the output of the model is **zero**. It means that both the body and tail stem cells are missing (**Fig. 18 & Eq.3**). Third, using Decision Tree: applying the Decision Tree rules to the head stem cell with no incoming signals, the output of DT is ‘**both**’ (**Fig. 20 left side**). Head stem cell identifies that both the body and tail stem cells are missing. The three approaches perform well for testing the missing stem cells in this case where stem cells are few. Automata and Decision Tree follow rules while the Neural Network calculates a numerical output value for making the final decision.

###### Replace missing smart stem cells

After identifying missing stem cells, the head stem cell reproduces a new body stem cell and a new tail stem cell and transfers the required information to each new cell from the shared information field. It means that the newly formed stem cells now have the shape information of the representative tissues.

###### Regenerate missing tissues

The new body smart stem cell produces a new rectangular body tissue while the new tail smart stem cell produces a new triangular tail tissue. The regeneration proceeds from a small body and tail shape attached to the head and the two new tissues grow concurrently, as shown in Fig. 24, maintaining the overall shape guided by the shape information *d, AR and n* (*number of corners*) accessed from the field and neighbourhood relations (as in **Fig. 5** and **Fig. 10**). After regeneration, the two levels of the self-repair system return to normal in that each smart stem cell returns to normal operation and maintains tissue entropy and the smart stem cell network gets re-established and resumes its normal operation where each stem cell receives the correct signals from the other stem cells.

**Fig. 24.**
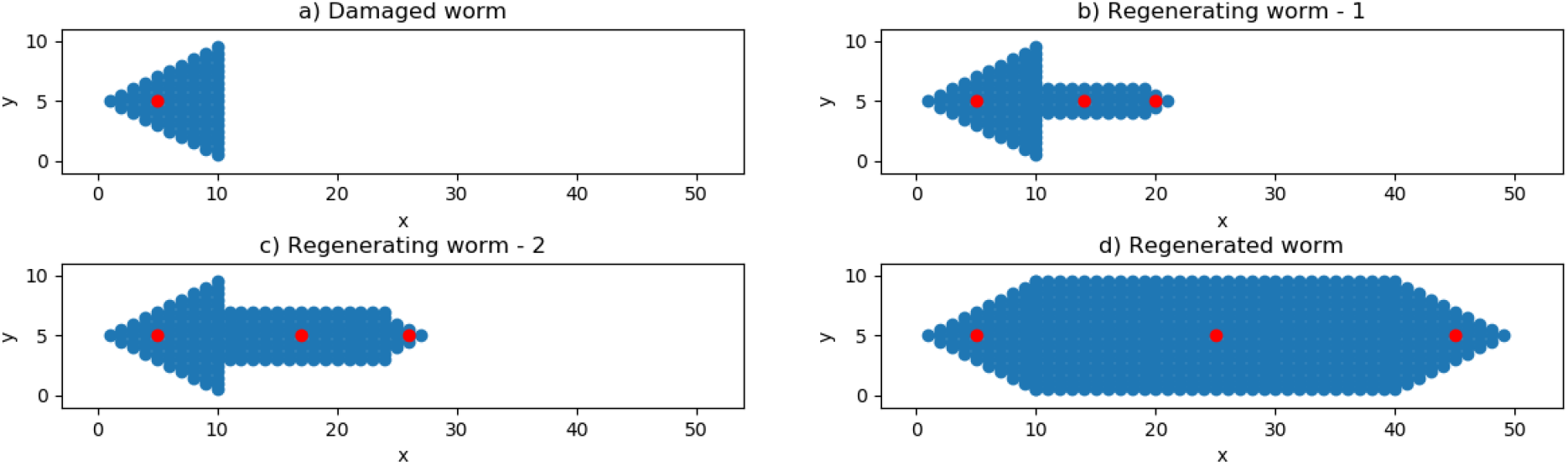
Progression of regeneration of body and tail from head

##### 5.2.1.2 (Body+Tail) regenerates head

Steps involved in head regeneration are similar to those in body and tail regeneration above. Body and Tail recognize the missing head stem cell using the same process as above. The three methods produced the same results efficiently. The body stem cell regenerates a new head stem cell and transfers the required shape information. Then, the new head stem cell starts producing a new triangular tissue and the regeneration continues until the sides become equal to *d* (as in Fig. 25). In this regeneration, pattern information for the triangular tissue (d, AR, n) is applied and the AANN and the neighbour relations get re-established. After regeneration, the two levels of the system return to normal operation.

**Fig. 25.**
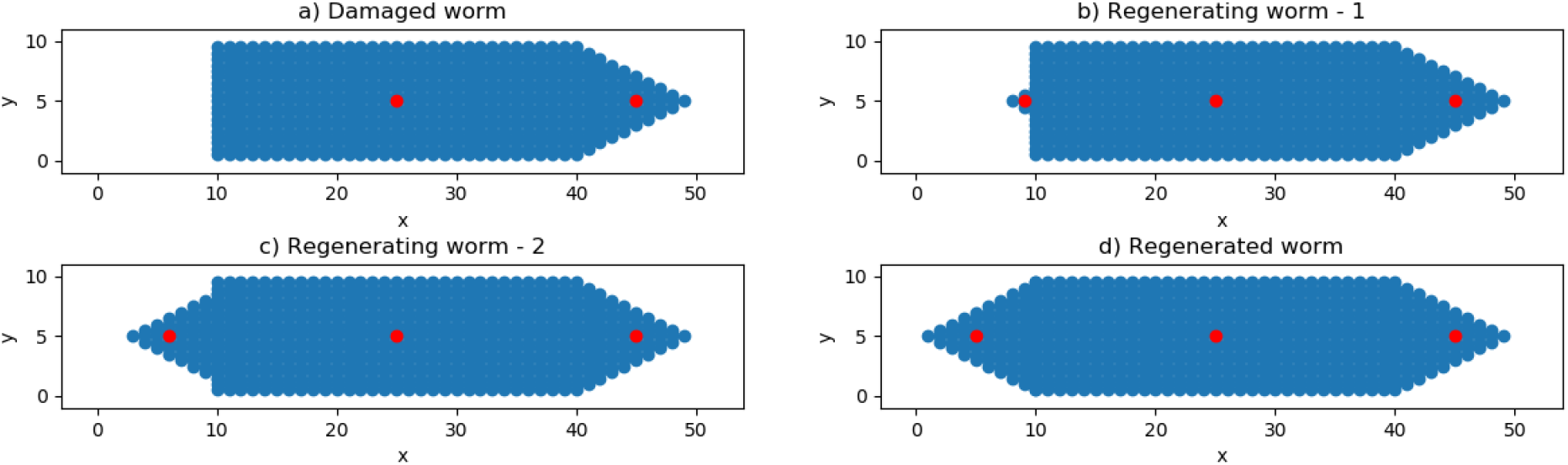
Progression of regeneration of head from Body+Tail

#### 5.2.2 Organism fragments into 2 parts encompassing damage to interior

In this case, the organism fragments into two parts – one small fragment of the body tissue containing the body stem cell is removed from the planarian leaving 2 damaged parts as in Fig. 26. Both parts will regenerate by activating the repair system differently.

**Fig. 26.**
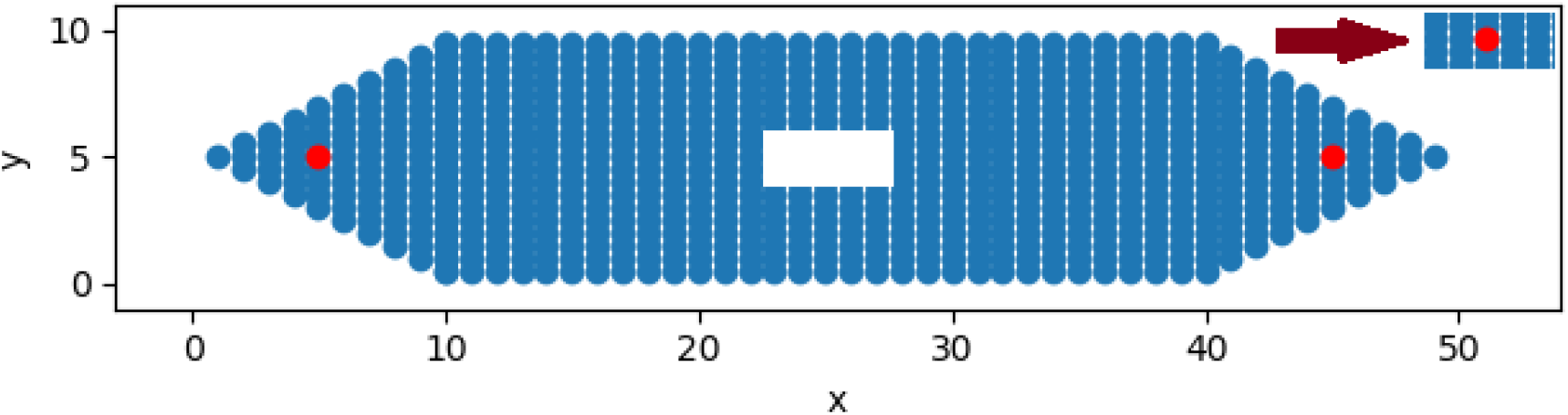
Interior damage to the worm removes a fragment of tissue with stem cell leaving two damaged parts

##### 5.2.2.1 The organism with missing stem cell regenerates into full organism

Regeneration of the larger part (body with missing stem cell) requires communication between the two repair sub-models (Levels 1 and 2). Damage signals activate stem cell repair sub-model and the process is as follows: (i) identify the missing stem cell (ii) replace missing stem cell, (iii) then activate only the (body) tissue repair model, (iv) activate AANN model to identify the damage border; (iv) regenerate. The repair process is described below.

###### (i) and (ii) Identify and replace missing stem cells

These two steps are similar to the previous case in 5.2.1a (i and ii) that applies the three approaches described in section 5.2. The system recognised that the body stem cell is missing. Either the Head or the Tail stem cell can produce the new body stem cell and transfer the required shape information to it.

###### (iii) Activate tissue repair sub-model (Level 1) – activate the AANN model to identify the damage border

As a large part of the body tissue remains in the organism, the new body stem cell does not need to regenerate the whole body tissue. Instead, it only needs to identify the border of the damage and repair the damage. It activates the AANN model in the tissue sub-model to find the damage border. This case does not require tapping into the information field for pattern information.

###### (iv) Regenerate missing tissues

After identifying the damage border, the new body stem cell migrates to the damage site (Fig. 27b) to regenerate the missing tissue until full recovery (Fig 27 c and d). After regeneration, the two levels of the system in the worm resumes normal operation.

**Fig. 27.**
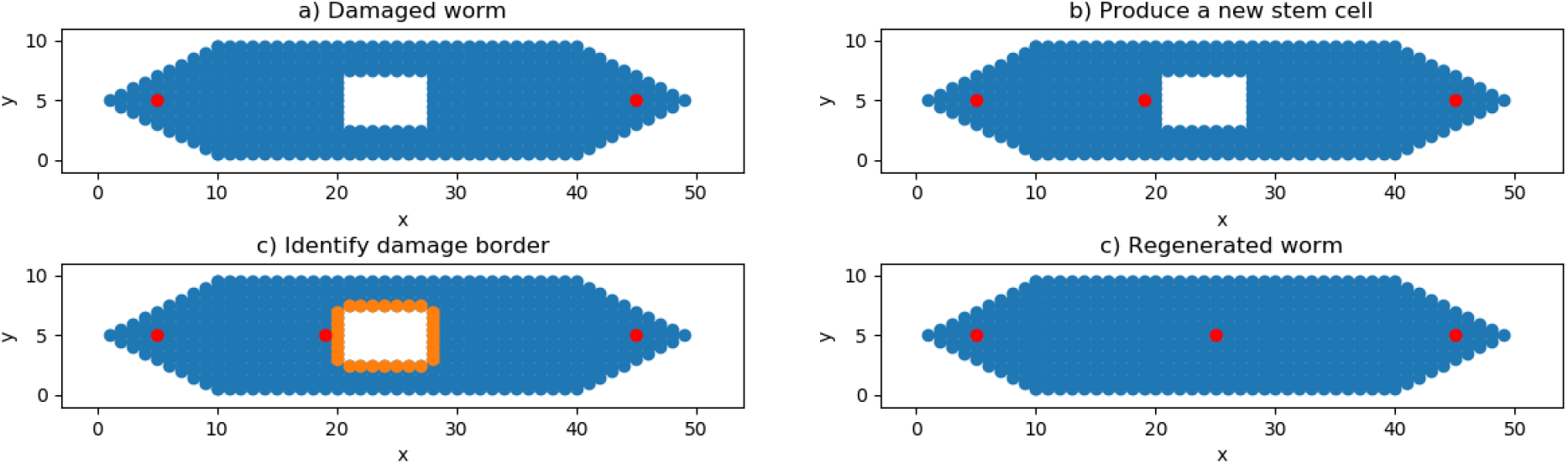
Progression of regeneration of missing interior tissue segments

##### 5.2.2.2 Small tissue fragment with body stem cell regenerates into full organism

The process is similar to the case in 5.2.1 above. As this damage leaves an open-ended boundary, the system accesses pattern information from the collective field. Damage signals activate the stem cell repair sub-model and the process is as follows: (i) identify missing stem cells, (ii) replace missing stem cells, (iii) regenerate missing tissues concurrently.

The first and second steps are similar to the previous cases in 5.2.1. The separated tissue with the body stem cell identifies and replaces the head and tail stem cells and transfers the required shape information to each new smart stem cell. The head stem cell produces a new triangular head tissue while the tail stem cell produces a new triangular tail tissue on the corresponding sides of the body tissue. However, how does the remaining stem cell know the direction to regenerate the head and tail tissue? Here, we assume that the stem cell network polarizes each stem cell (+,-) depending on their relative position indicated by the cell identifier (1, 2, and 3 in our example) where “+” and “-” show the direction of the head and tail, respectively, as shown in Fig. 28. A stem cell with a smaller identifier relative to a neighbour cell indicate “+” direction. This property of the stem cells helps them identify and grow in the correct direction as in the original. The three smart stem cells self-organize to become a new small worm (Fig. 29b) and then the three parts grow concurrently while maintaining the overall shape. Regeneration continues as shown in Fig. 29b-d until the sides of the head and body become equal to *d* and the body reaches aspect ratio of 3. After regeneration, the two-level system in the organism resumes normal operation.

**Fig. 28.**
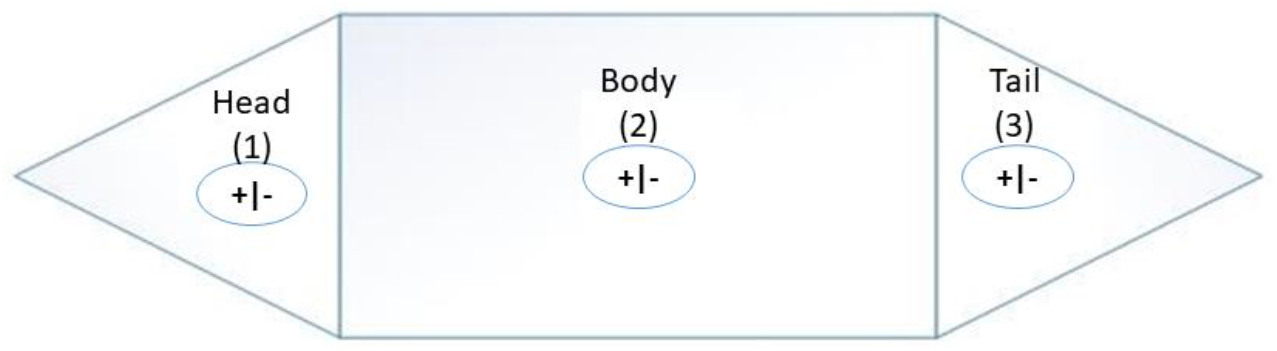
Smart stem cell polarities

**Fig. 29.**
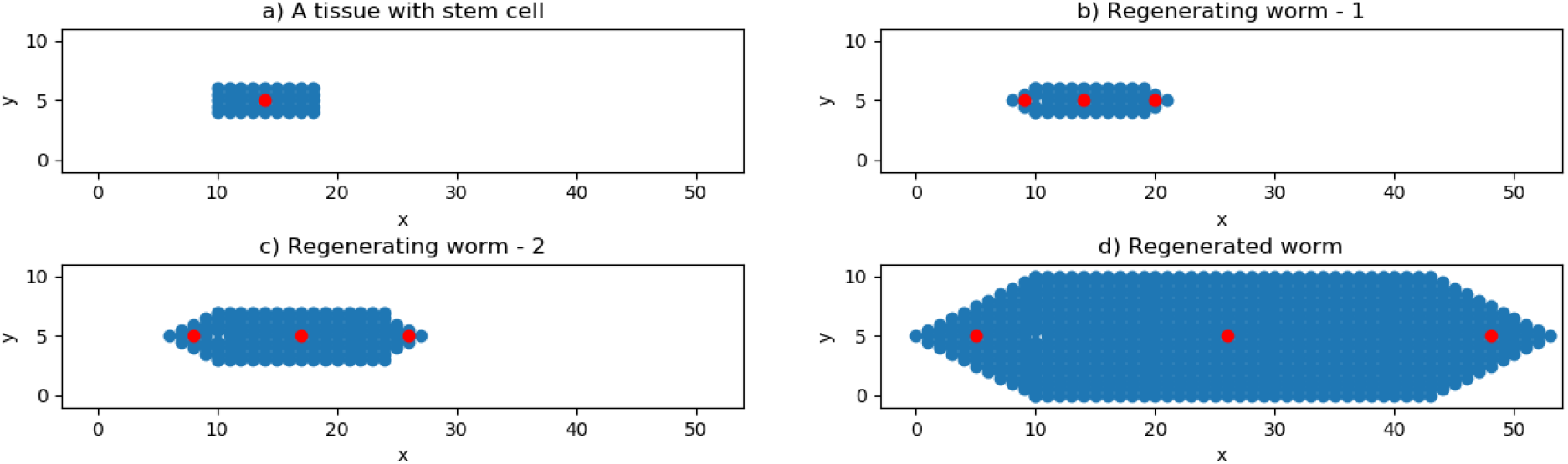
Steps in the regeneration of fragmented body tissue with stem cell into a complete organism

## 6. Discussion

We proposed a whole system regeneration model with minimum computation based on a combined operation of tissue and smart stem cell self-repair systems to completely and accurately recover from any damage anywhere in the system. We demonstrated its efficacy in recovering from a number of small, large and severe damages. The system represents tissues with smart stem cells that are aware of their pattern integrity maintained through a measure of entropy and a neural network of differentiated cells that maintains the pattern. Each smart stem cell keeps a small amount of pattern information in its virtual field that is shared with other smart stem cells through a collective field that is accessed by the stem cell repair network. The two networks seamlessly collaborate to accomplish complete regeneration.

Our model makes a number of conceptual and algorithmic contributions to advancing the capability of regeneration models and improving the understanding of regeneration. In this regard, we make the following comparison of the model to the previous models of regeneration. In (Bessonov et al., 2015) model, all cells send signals to each other, which is not computationally efficient and effective due to a great number of communications between cells. In our model, only the stem cell communicates with all other cells in a limited fashion while other cells only communicate with their (≤ 4) neighbours. Their model requires extensive computation for calculating the total received signals for each cell while in our model, only the stem cells need to calculate the entropy when they monitor the system. AANN and stem cell network do limited computation as and when needed because the AANN only activates a small region of its space where the damage is as guided by the stem cell.

Model of (Tosenberger et al., 2015) described the development and regeneration of a structure where the global submodel presents the restoration of tissue centres (e.g., stem cells), while the local model produces tissues around the stem cell. Our model is similar and different to (Tosenberger et al., 2015) model in some aspects. First, a stem cell network in our model is similar to their global model. They alter the position of stem cells and make them return to the initial positions, but it cannot replace new stem cells. In our model, the stem cell network can monitor the state, detect damage and produce new stem cells to replace missing stem cells. Second, they proposed different types of signals that provide information to restore the exact initial configuration of the cell structure when the position of cells are altered except for the displacement of the stem cells in diametrically opposite directions. In our system, stem cells produce the same type of signal and stem cell network can regenerate stem cells in the exact locations, such as head, body and tail. In our model, the regeneration of individual parts is different from the local model of Tosenberger in some aspects. We use entropy to detect general damage region, AANN to detect the exact location and the border of damage and neighbourhood rules for regenerating the pattern without creating even a single additional or redundant cell, whereas, they assumed that a stem cell can produce and maintain a circular tissue based on an assumption that the stem cell has a survival region, and a differentiated cell located beyond this region dies as it receives a signal smaller than a threshold. Moreover, the local model in (Tosenberger et al., 2015) only applied to a simple shape like a circular tissue while our model works successfully on other shapes such as triangular and rectangular shapes.

(De et al., 2017) model involves two levels of interaction between a neural skeleton and non-neural cells that coordinate tissue growth using biologically realistic variables and concepts. However, it does not recognise damage, and generates and kills many cells before reaching a partially recovered form. In our system, minimum pattern information is stored in the stem cell virtual fields that are accessed only in extreme damage cases; for the majority of damages, it uses cues from the remaining pattern for regeneration. A stem cell monitors the tissue integrity through entropy and the tissue maintains neighbourhood and border rules. Our model also detects damage and recovers the complete form using simpler and efficient computations. As long as a single smart stem cell exists, it can regenerate the whole organism simply and easily, while in De et al. model, the percentage of accuracy of regeneration is decreased when the injury is increased.

The agent-based models (Ferreira et al., 2016; Ferreira et al., 2017a; Ferreira et al., 2017b) need to store too much information (paths of packets) in stem cells as well as too much communication between cells in discovering states. The information our model carries for regeneration is high-level and generic – minimum information on shape and entropy and neighbourhood rules. These other models are also limited when it comes to regenerating large sized organisms due to the inability of the packets to go too far or signals decaying with distance. By contrast, our model allows a correct regeneration for an organism of any size.

Although our model emulates some important aspects of regeneration in biology, it still has some limitations that need to be considered. First, there are only three stem cells in the proposed model that seem unrealistic for an organism. In the next stage, we plan to increase the number of stem cells to about 20% of the total cells (like in the planarian). Second, the pattern of the organism is simple with head, body and tail. We will also extend the organism to a more complex form with two eyes and pharynx. Finally, the three methods used for stem cell repair network, Automata, Neural Networks and Decision Trees, perform with similar efficiency in the damage cases tested in this study. However, when the stem cell population in the network becomes larger as in a model of the real planarian, the Automata may not be useful while the neural network may become more efficient. The Decision Trees may become helpful for a larger population of stem cells when each type of stem cell (e.g., head/body/tail stem cells) has a property indicated by the cell identifier. We plan to further explore the efficacy of these methods in the next planarian regeneration model. Further, our model is a framework representing an abstract concept for complete regeneration without explicitly incorporating the underlying biological processes. However, the model can be expanded incorporating these processes, such as cell cycle and bioelectric patterning related mechanisms, that operate under the developed framework. We hope to incorporate some of these biophysical variables in the next stage of model development.

## 7. Conclusions

In this study, we modelled computational dynamics that enable regeneration in a simulated biological system. We developed a comprehensive conceptual framework for modeling autonomous complete regeneration systems that mimicked the observed patterns and capacity of regeneration in organisms like planaria. The proposed conceptual model is very robust in returning the system to the normal state after any damage. Specifically, our model is consistent with observations of planarian regeneration, including complete and accurate regeneration of any part or whole organism from any damage. The minimum requirement for regeneration is that only a single stem cell remains after damage. The model also provides a different perspective on the role of local (short-range) interactions (between differentiated cells), long-range interactions (between smart stem cells) and potential information fields during regeneration that presents a hypothetical paradigm for understanding the regeneration processes. It would be of interest if biological experiments could reveal these short and long-range communications and any proxy measures for the existence of information fields. Further, our model could be the basis for self-repair systems in synthetic biology – bioengineered and fully artificial robots. The success of the model suggests the possibility for a biological system to autonomously sense, detect and repair damages completely. In the next stage, the model will be extended by investing the neurons in the AANN with greater functionality in terms of further information processing, temporal learning and reasoning; improving cognition capabilities to enhance collective intelligence and decision-making, and incorporating properties of living cells such as bioelectric signals. The aim will be to exploit the model to provide insight into general self-recognising, self-organising and self-stabilising cell communities that are intelligent decision-making systems with increasingly greater resemblance to living cell systems.

## Acknowledgments

Authors gratefully acknowledge support of the following: TNM – Doctoral Scholarship from VIED, Vietnam; S.S.- LURF research Fund (New Zealand); M.L.- DARPA (#HR0011-18-2-0022), the Allen Discovery Center award from the Paul G Allen Frontiers Group, and the Templeton World Charity Foundation (TWCF0089/AB55 and TWCF0140).

